# A decision-based criterion and a resolution threshold for ranking genomic prediction models within trials

**DOI:** 10.64898/2026.10.03.756375

**Authors:** Guohao Lv, Renzhe Zheng, Lichuan Gu

**Affiliations:** School of Artificial Intelligence, Anhui Agricultural University, 130 Changjiang West Road, Hefei 230036, Anhui, China; Anhui Province Key Laboratory of Intelligent Agricultural Technology and Equipment, Anhui Agricultural University, 130 Changjiang West Road, Hefei 230036, Anhui, China

**Keywords:** genomic selection, genotype-by-environment interaction, evaluation metrics, selection response, prediction competition

## Abstract

Breeding programmes choose genomic prediction models by accuracy rankings, and the metric behind a ranking decides which model wins. Selecting the best genotypes within a trial depends only on the order of the predictions, so a metric that guides the choice must be unchanged by any order-preserving change of the predictions and respond when their order is reversed. Of the 22 official metrics of the Genomes to Fields maize competition, one meets both conditions, the within-environment rank correlation; the within-environment Pearson correlation is protected against rescaling only; and twenty fail, including the root mean squared error that decided the 2022 ranking. Switching between the competition’s two official metrics reverses 27 % of team pairs, and an exact decomposition shows why pooled metrics mislead: in three constructed panels, 94–95 % of the attainable square of pooled correlation is carried by the environment means. The accuracy difference a trial can resolve falls as about 3/√*N* with its *N* genotype–environment cells: at least 0.03 at the competition’s design and 0.04–0.12 once calibrated on real panels, against 0.003 between the first two teams; tests that two correlations differ declare far smaller differences, which do not reliably change selection. In soybean, where the two most accurate methods stand clear of most others by more than the trial resolves, ranking by either within-environment correlation selects better material, 98–99 % of the attainable selection gain in held-out environments against 13–34 % for error-magnitude rankings; in common bean and a Chinese maize population, where the leading methods lie within the resolvable gap, all rules pick material of similar value. GPverdict runs these checks on any programme’s cross-validated predictions.

## 1. Introduction

Genomic prediction has become the organising method of modern variety development. Since Meuwissen et al. [1] showed that dense marker maps could predict total genetic value, the approach has moved from proposal [2] to routine practice across the major crops [3], and is credited with much of the acceleration in genetic gain that breeding programmes now report [4]. Its progress is tracked almost universally by a correlation or an error: how closely predicted values track observed ones [5,6].

Multi-environment prediction is where the method is hardest and most actively developed. Genotype-by-environment interaction has been absorbed into the prediction model through pedigree and marker covariance structures [7], reaction norms on environmental covariates [8,9], marker-by-environment terms [10], envirotyping [11,12], crop growth models [13] and deep learning [14,15]. Public competitions now rank dozens of independent teams on a single published number [16].

A breeder, however, does not consume a prediction as a number. They consume it as a decision — which genotypes to advance, and which to discard. That decision compares genotypes only against others grown in the same environment, and it depends only on their order. Any transformation that preserves that order within each field — a shift, a rescaling, or a non-linear distortion — produces the identical decision; a prediction that is systematically biased but correctly ordered costs nothing. An accuracy metric that responds to such a transformation is responding to something the decision cannot see.

That evaluation metrics can disagree is well documented. In genomic selection the best method depends on whether it is scored by Pearson correlation or by a ranking measure [17], and in the G2F competition a team moved from 15th to 4th place when the metric changed [16]; competition winners in biomedical image analysis change with the metric variant [18,19], and the top-ranked winter wheat genotypes in variety trials depend on the model that ranks them [20]. Genomic prediction has debated how accuracy should be estimated [21,22] and models compared by cross-validation [23], whether Pearson correlation alone suffices [24,25], and whether tail-selected measures serve better [26–28]; decision theory has been applied to the choice of genotypes [29], not of metrics. Recent work shows predictive ability and accuracy diverging between model classes [30], cross-validated accuracy overstating what genomic selection delivers [31], a widely adopted model class not earning its reported advantage [32], pooled accuracy masking weak within-environment accuracy [33], and benchmarks confounded by field variability [34].

The work closest to ours is that of Eckhoff et al. [35], who showed in hybrid rye that losses and metrics pooled across environments are dominated by environmental main effects, trained and scored models within environments, and argued that the mean squared error of differences (MSED), because it also registers shrinkage, is a more complete target than the within-environment Pearson correlation. Buntaran et al. [36] argued that predictive accuracy does not answer the breeder’s question and simulated instead the probability that the selected set contains the truly best genotypes, and Rio et al. [37] predicted the reliability of genomic predictions to optimise trial design. Within-environment rank correlation, MSED and its location-wise form MSEPD, the coincidence index and NDCG at *k* are all in current use for multi-environment and selection-oriented evaluation [28,38,39], usually side by side and without a rule for choosing among them when they disagree. None states which of the accuracy metrics already used to rank genomic prediction methods are *admissible* for a given selection decision, says how large a reported improvement must be before the more accurate method reliably wins it, or tests whether the metric so chosen leads to better material in environments it has not seen.

Here we answer all three for within-environment truncation selection — one of the decisions by which genotypes are advanced, taken on their performance in a single trial. We define admissibility by the transformations that leave the decision unchanged, derive the accuracy difference a trial can resolve from its genotype–environment cells, test the metrics against the material each one’s choice of method selects in held-out environments, and show exactly why pooled metrics reward the environment means. The Genomes to Fields (G2F) maize competition provides a direct test, because its organisers changed the official ranking metric between editions; three datasets in three further species test whether the answers generalise; and an open tool, GPverdict, returns the same answers for any programme’s predictions.

## 2. Materials and methods

### 2.1 The admissibility criterion

The decision considered is truncation selection within an environment: rank the genotypes grown in environment *e* by their predictions and advance the top *k*, for whatever *k* the programme chooses. For any strictly increasing transform *g*, replacing the predictions ŷ*ₑ* by *g*(ŷ*ₑ*) leaves the advanced set unchanged for every *k*; reversing them, ŷ*ₑ* → −ŷ*ₑ*, turns the best selection into the worst. A metric can represent the decision only if it is invariant under the first class and changes under the second (Fig. 1A). We call a metric meeting both conditions **admissible** for the decision (Tier I), in the sense of measurement theory rather than of statistical decision theory.

**Fig. 1.**
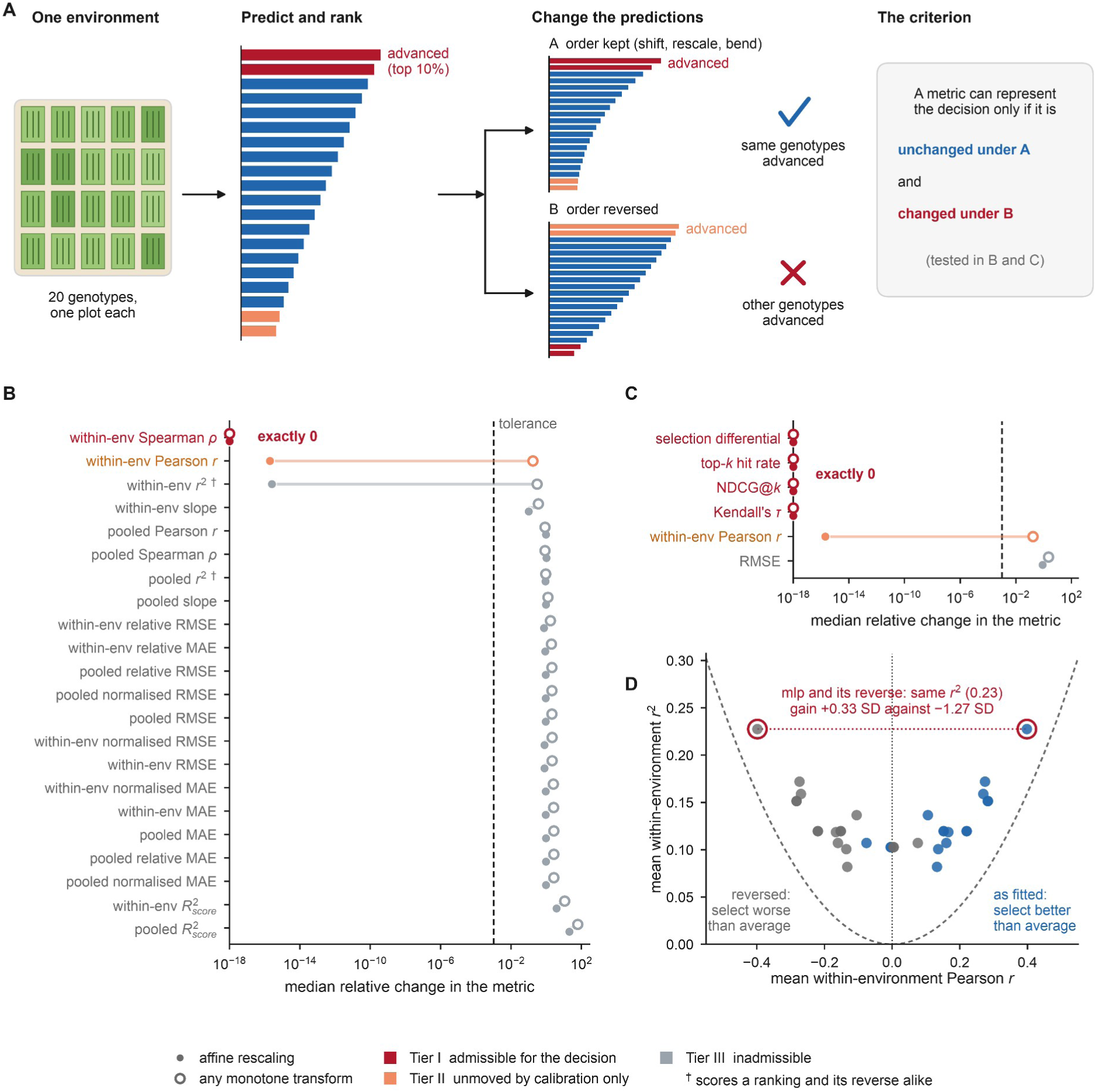
One official metric is admissible for within-environment selection, and one more is protected against calibration only. (A) The criterion: an order-preserving change of the predictions (i) advances the same genotypes, whereas reversing their order (ii) advances others; an admissible metric is unchanged under (i) and changed under (ii). Bars are illustrative. (B) Median relative change of the 22 official metrics under within-environment affine rescaling (filled) and random monotone transforms (open) of five verified G2F submissions. Dashed line, the 10⁻³ tolerance; colour, tier; †, squared correlations, which score a ranking and its reverse alike. Supplementary Table S2 gives the official metric names. (C) The same tests for decision-oriented measures in maize; RMSE moves by 0.81 under rescaling and 2.15 under the monotone transform. (D) The seventeen spring wheat base methods as fitted (blue) and reversed within every environment (grey) receive the same *r*². Axes are means over environments; dashed, the square of *r*, which the points exceed because *r* varies among environments.

#### Proposition

*A metric M*(ŷ, *y*) *is invariant under all strictly increasing within-environment transformations of* ŷ *if and only if it depends on* ŷ *only through the within-environment rank pattern of* ŷ, *ties included*.

The proof is in Supplementary Section S1; ties straddling the *k*-th place leave the selected set itself non-unique (Supplementary Section S10).

The rank pattern is thus a maximal invariant of the transformation group — the standard characterisation of invariant functions. The class follows from what is fixed in advance: a ranking meant to serve any selection fraction must support the decision over all *k*, the case analysed here; when a programme fixes its fraction instead, the admissible metrics are those of the selected set, such as the realised selection differential or the top-*k* hit rate at that fraction. Both classes exclude every metric that reads the scale of the predictions.

Restricting the class to per-environment affine maps ŷ*ₑ* → *a*ₑŷ*ₑ* + *b*ₑ with *a*ₑ > 0 gives a weaker property. A metric invariant under it cannot be moved by calibration differences — location and scale within each environment — but can be moved by non-linear order-preserving distortions that leave every selection unchanged. Order-sensitive metrics with only this property form Tier II; all others are Tier III. Membership follows from the definitions — a within-environment rank correlation is invariant by construction, a Pearson correlation affine-invariant, a squared correlation unchanged by reversal — so the tests below verify the implementation and measure how far the failing metrics move while every selection decision stays fixed.

This is the meaningfulness criterion of measurement theory applied to evaluation rather than to data: a statement is meaningful only if its truth survives the transformations the underlying scale admits [40,41]. Here the decision, rather than the measuring instrument, fixes the transformation class. The same reasoning governs which scoring functions rank forecasts consistently [42] and on which scales information-retrieval measures live [43], and it explains why the error-magnitude family fails: RMSE is consistent for the conditional mean [44], which a within-environment ranking neither determines nor needs.

Admissibility is defined relative to a decision. Selecting on predicted genotype means across environments admits the metrics that depend on the predictions only through the ranking of those means; selecting on an index of traits, or with genotypes weighted by economic value, fixes its own class in the same way. The criterion constrains how a metric uses the predictions, not the phenotypes: a metric that keeps observed values on their original scale, as the realised selection differential does, remains admissible. Within the admissible class, a metric is then chosen for its weighting of the selected tail and its statistical efficiency.

### 2.2 Datasets

Four multi-environment datasets spanning four species and two trait classes were analysed (Table 1).

**Table 1.** Datasets and principal results. Environments and genotypes per environment describe each dataset’s evaluation set, and genotypes per environment is the median over its environments; the soybean analyses use part of that set (‡). Rank ranges and interval widths are fractions of the board. ᵃ Thirty teams on the leaderboard; five verified submission files from three teams carry per-genotype predictions, too few to estimate reliability. ᵇ Gaussian value at the competition’s design; in parentheses, the range after scaling by the three panels’ observed-to-Gaussian ratios. The threshold’s upper confidence limit lies beyond the observed range in all three panels. ‡ The forward-year evaluation subset of the soybean series; the soybean analyses use the 169 of its environments that carry at least 25 genotypes, with 1,161 genotypes, and the design match of Supplementary Table S10 uses the median over the method–environment cells that carry at least 25 genotypes, which in soybean is 61.

| Measure | maize | common bean | spring wheat | soybean |
| --- | --- | --- | --- | --- |
| Dataset / trait | G2F 2022 / yield | VEF / yield | URSN / FHB resistance | NUST / yield |
| Methods compared (base methods) | 30 teams<sup>a</sup> | 52 (17) | 52 (17) | 66 (26) |
| Environments / genotypes per environment | 23 / 445 | 10 / 344 | 109 / 16 | 184 / 55‡ |
| Markers | not used | 5,820 | 2,302 | 5,158 |
| Official metrics in Tier I / II / III | 1 / 1 / 20 | 1 / 1 / 20 | 1 / 1 / 20 | 1 / 1 / 20 |
| Reversal between the two official metrics | 26.9 % | 39.7 % | 47.2 % | 34.7 % |
| 95 % CI (teams or method families resampled) | 18.0–37.1 | 31.4–44.9 | 39.1–53.6 | 26.0–42.5 |
| Reversal without the added variants [95 % CI] | — | 16.2 % [5.7, 31.0] | 25.0 % [12.7, 40.5] | 19.8 % [10.8, 28.8] |
| Median rank range across the official metrics | 40 % | 50 % | 53 % | 52 % |
| Median 95 % rank-interval width | 57 % | 75 % | 85 % | 79 % |
| Outcome reliability | —<sup>a</sup> | 0.869 | 0.647 | 0.786 |
| Threshold $\Delta r^*$ at $f=0.10$ [lower limit] | 0.028 (0.04–0.12)<sup>b</sup> | 0.066 [0.024] | 0.213 [0.056] | 0.116 [0.038] |

#### Maize (*Zea mays* L.)

G2F 2022 G×E prediction competition; data release [45] and results from Washburn et al. [16]. Thirty teams were scored on a 2022 test set of 23 environments, with 22 metrics per submission published in Supplemental Table S1 of that paper: 124 submissions, 116 of them carrying all 22 metrics. As on the official leaderboard, a team is represented by its best submission under the 2022 official metric (four alternatives in Supplementary Table S21). Five submission files recovered from the team repositories reproduce the official values to within 5 × 10⁻⁷ and are used wherever per-genotype predictions are required (Supplementary Section S19).

#### Common bean (*Phaseolus vulgaris* L.)

VEF Andean elite panel of Keller et al. [46], data deposit [47]; 467 lines, 5,820 SNPs, 10 trials in Colombia under drought, irrigated and phosphorus regimes, seed yield.

#### Spring wheat (*Triticum aestivum* L.)

Uniform Regional Scab Nursery 1995–2024 [48], data deposit [49]; 384 lines, 2,302 markers, 109 environments, Fusarium head blight disease score (DIS, 0–100) analysed as resistance = 100 − DIS.

#### Soybean (*Glycine max* (L.) Merr.)

Northern Uniform Soybean Tests [50]; 2,513 genotypes, 626 environments, 2003–2020, 5,158 SNPs. Yield was predicted forward in time, training on all earlier years and testing on each of 2016–2020 in turn, so the series contributes 184 environments and 1,162 genotypes. The analyses below use the 169 of those environments that carry at least 25 genotypes, which leaves 1,161 genotypes (Table 1, footnote ‡).

Population structure is described in Supplementary Section S23 (Fig. S4). Rice and wheat panels built from two further public datasets were withdrawn because their methods ranked genotypes within environments near chance (Supplementary Section S16).

### 2.3 Metric panel and prediction panels

As in the competitions, accuracy here means agreement between predictions and observed phenotypes — predictive ability in the terminology of ref. [30]. For every prediction vector we computed the 22 metrics the G2F organisers reported [16], both pooled across all genotype–environment cells and within environment and then averaged (the mean_ family): RMSE, MAE, their normalised and relative forms, *R*²_score, Pearson *r*, *r*², Spearman *ρ*, and the slope of observed on predicted. Breeding software computes per environment where it groups at all [51]; general-purpose libraries and genomic-prediction panels [52] score pooled vectors. Environments were required to carry at least 25 genotypes, except in the spring wheat nursery, whose design gives a median of 16 lines per environment: a floor of 25 would leave 16 of its 109 environments, so a floor of 12 was used there, which keeps all 109 environments. The maize submissions entering the invariance tests used a floor of 20. Metrics were oriented so that higher is better; slopes were scored as −|slope − 1|.

The other three datasets have no competition, so a panel of 17–26 base methods was constructed for each (Supplementary Table S20): ridge regression on an environment index and marker principal components at four dimensionalities crossed with two penalties [53], random forests, histogram gradient boosting, *k*-nearest neighbours, a multilayer perceptron [14,15], unweighted ensembles and a marker-only ridge, together with the two mixed models most used in multi-environment genomic prediction — a PC-kernel GBLUP on the same principal components and its marker × environment extension [10], reduced-rank models on the feature space every method in these panels shares so that the comparison isolates the learner, fitted by restricted maximum likelihood; the soybean panel adds a genomic Finlay–Wilkinson reaction norm [54]. Cross-validation was five-fold leave-genotypes-out for bean and spring wheat and forward-year for soybean, the CV1 and forward-prediction designs of the multi-environment literature [7]; the composition of the training set is known to matter to the accuracy obtained [55]. Learners were run at conventional settings rather than tuned, and no test phenotype enters a prediction (Supplementary Sections S12 and S19).

Panels fitted by least squares are far more homogeneous in calibration than competition entries (Supplementary Section S19), so each was augmented with shrunk (×0.25 about the mean), inflated (×3), biased (+0.8 SD), noise-injected and partially rank-shuffled variants of seven of its members (eight in soybean). The first three are affine copies that rank genotypes exactly as their parent does, so pairs tied in accuracy or in outcome are dropped wherever pairs are compared; every analysis is also reported on the unaugmented panels.

### 2.4 Statistical procedures

#### Invariance and order tests

A metric is admissible if a random strictly increasing within-environment map applied to the predictions moves its median relative change by less than 10⁻³ and reversing every environment moves it by at least 10⁻³; the partition does not depend on that tolerance (Supplementary Section S2). *Reversal rate.* A reversal is Δ₁ · Δ₂ < 0 in the oriented score differences, so the rate is the normalised Kendall distance between the two rankings [56]; pairs tied on either metric are excluded, and intervals are cluster bootstraps resampling teams in maize and method families in the panels (Supplementary Section S4). *Rank intervals.* Each metric is treated as a task, each method’s interval is centred on its median rank with a half-width common to all methods, and the interval is descriptive rather than conformal because it is fitted and evaluated on the same cells [57] (Supplementary Section S9; Supplementary Table S6). *Realised selection differential.* For each method and environment the top *k* genotypes by prediction were selected, with *k* = *fn* rounded to the nearest integer, *n* the number of genotypes in the environment and *f* = 0.05, 0.10 or 0.20; the realised selection differential is the mean observed value of the selected genotypes less the environment mean, in phenotypic standard deviations.

#### Threshold

For every pair of base methods sharing at least eight environments, Δ*r* is the mean difference in within-environment Pearson correlation and Δ*g* the mean difference in realised selection differential at *f* = 0.10, oriented so that Δ*r* ≥ 0; tied pairs are dropped. Relabelling the two methods leaves their joint distribution unchanged, so the probability that the more accurate method also has the larger Δ*g* is exactly 1/2 at Δ*r* = 0. We therefore fit *P(agree | Δr) = 1/[1 + exp(−β Δr)]* by maximum likelihood and define the threshold Δ*r*\* = logit(0.95)/β; 95 % intervals resample methods and environments together (two-way cluster bootstrap, 500 replicates). The same estimator applied to simulated Gaussian panels gives a design-based reference; for maize, whose five verified submissions give ten pairs, the Gaussian value at the competition’s design was scaled by the three panels’ ratios of observed to Gaussian thresholds (Supplementary Section S19; Supplementary Table S10).

#### Outcome test

Environments were split at random into halves; each of six scoring rules (mean_pearson_r, mean_spearman_r, NDCG@10 %, mean_RMSE, mean_MAE and mean_r2_score) ranked all methods on half A and the realised selection differential of its top-ranked method, selecting the top 10 % of genotypes, was measured on half B, over 400 splits, and expressed as the fraction of the distance from an average method to the best method on half B. Datasets whose split-half reliability falls below 0.5 are reported but not compared, and the test was repeated for a threshold decision and for selection on genotype means (Supplementary Sections S8 and S15).

#### Metrics from recent studies

The mean squared error of differences (MSED) [35], its location-wise form MSEPD [38], the coincidence index [39,58] and NDCG at *k* [28] were put through the same tests (Supplementary Table S12).

#### Tests of correlations

For the same pairs, the accuracy difference was also tested by the test of Meng et al. [59] and by a *t* test over environments (Supplementary Section S22).

#### Pooled and within-environment components

Pooled Pearson correlation and pooled mean squared error were split exactly into a part carried by the predicted environment means and a within-environment part (Supplementary Section S20).

#### Supplementary analyses

Sections S1–S23 (Tables S1–S32; Figs. S1–S5) give the proof of the criterion and every supplementary analysis cited here, including a noise null (disagreement between two metrics is 3.0–7.0 times one metric’s sampling variability; Supplementary Table S3), the shortfall of realised selection below its projection *i*ₖ · *r* (selection intensity times accuracy) in the verified maize submissions (Section S19), the ceiling that plot error sets on a within-environment correlation (Section S6), the withdrawn panels and GPverdict’s validation.

#### Software

Analyses were run in Python 3.11 and 3.12 with NumPy 2.4.6, SciPy 1.17.1, pandas 2.3.3, matplotlib 3.11.1 and scikit-learn 1.9.1.

### 2.5 GPverdict

The criterion, the resolution threshold and the outcome test are implemented in GPverdict [60], an open Python package with a command-line interface and a web version that runs in the browser without uploading data (Data and code availability). From a table of environment, genotype, observed value, prediction and method, it reports the empirical tier of eleven metrics on the user’s own predictions; the methods ranked by the within-environment rank and Pearson correlations, with 95 % rank intervals from resampled environments; the methods within *k*/√*N* of the best on either correlation, with *k* interpolated from the Gaussian design grid (Section 3.3) and *N* the genotype–environment cells; the method an RMSE ranking would choose; the outcome test, splitting genotypes within environments when there are fewer than eight environments; and the cells needed to resolve a given gain. It reproduces the spring-wheat results reported here exactly (Supplementary Section S18).

## 3. Results

### 3.1 Which metrics can represent the decision: one of twenty-two

Of the 22 official metrics, **exactly one is admissible**: mean_spearman_r is exactly invariant under the monotone test and changes sign when the ranking is reversed. mean_pearson_r is invariant under the affine test (drift 2 × 10⁻¹⁶) but moves by 0.173 under the monotone test, and is Tier II. mean_r2_pearson is also affine-invariant, but squaring discards the sign, so it awards a method and its precise reverse the same score: in spring wheat, 0.23 to both, although the first selects a top decile 0.33 phenotypic standard deviations above its environment mean and the second one 1.27 below (Fig. 1D). A metric that cannot separate a ranking from its reverse cannot rank methods. It and pooled *r*² therefore join the error-magnitude metrics in Tier III; the fourteen error-magnitude metrics move by 1.7 to 61 under the monotone test and by 0.75 to 21 under the affine test (Fig. 1B; Supplementary Table S2). Aggregation matters as much as the quantity: pooled Spearman, computed across environments rather than within them, moves by 0.85 under the monotone test.

**The metric that ranked the 2022 competition, mean per-environment RMSE, is in Tier III. The metric adopted for the 2024 edition, mean within-environment Pearson *r*, is in Tier II**: differences in calibration between methods cannot move it, but non-linear distortions that preserve every ranking can.

Because the partition follows from the definitions, it recurs in every dataset — one, one and twenty in maize, common bean, spring wheat, soybean and a fifth dataset, Chinese maize (Table 1; Supplementary Section S11) — confirming the implementation. It is also a partition of rankings, not only of values (Supplementary Table S14).

The criterion also admits, and gives a formal basis to the decision-oriented measures proposed elsewhere: the realised selection differential, whose normalised form is the relative efficiency of Ornella et al. [26], the top-*k* hit rate (and hence the coincidence index of plant breeding [58]), NDCG at *k* [17,61] and Kendall’s *τ* are invariant under both tests in maize, common bean and soybean (drift at most 5 × 10⁻⁶), while RMSE on the same predictions moves by 0.81–1.40 under the affine test and 2.15–3.66 under the monotone test (Fig. 1C shows maize).

Of the metrics used in recent studies (Supplementary Table S12), the coincidence index and NDCG at 10 % and 20 % are Tier I at a fixed fraction in all four datasets. MSED and MSEPD compare genotypes within an environment, so its location cancels, but they respond to rescaling: under the affine test rooted MSED moves by 0.6–6.0 % and MSEPD by 1.0–11.6 %, against 91–172 % for RMSE on the same predictions, and both are Tier III.

### 3.2 How far the choice moves a ranking: a quarter of team pairs, twelve of thirty places

The G2F organisers ranked the 2022 competition by per-environment RMSE averaged across environments; the 2024 edition [62] ranks entries by the mean within-environment Pearson correlation on its public leaderboard [63]. Between those two official metrics, **117 of 435 team pairs reverse order — 26.9 %, team-bootstrap 95 % CI [18.0 %, 37.1 %]**, and 15 of the 45 pairs among the 2022 top ten (Fig. 2). Median rank change is 4.5 places, the largest fifteen, and the top-three set differs between the two metrics. Across all 231 pairs of the 22 metrics the median reversal rate is 11.7 %, and 129 pairs reverse more than 5 % of team pairs under Benjamini–Hochberg control at FDR 5 % (84 at 1 %); for the two official metrics the adjusted *P* is 2.8 × 10⁻⁵ (Supplementary Section S4).

**Fig. 2.**
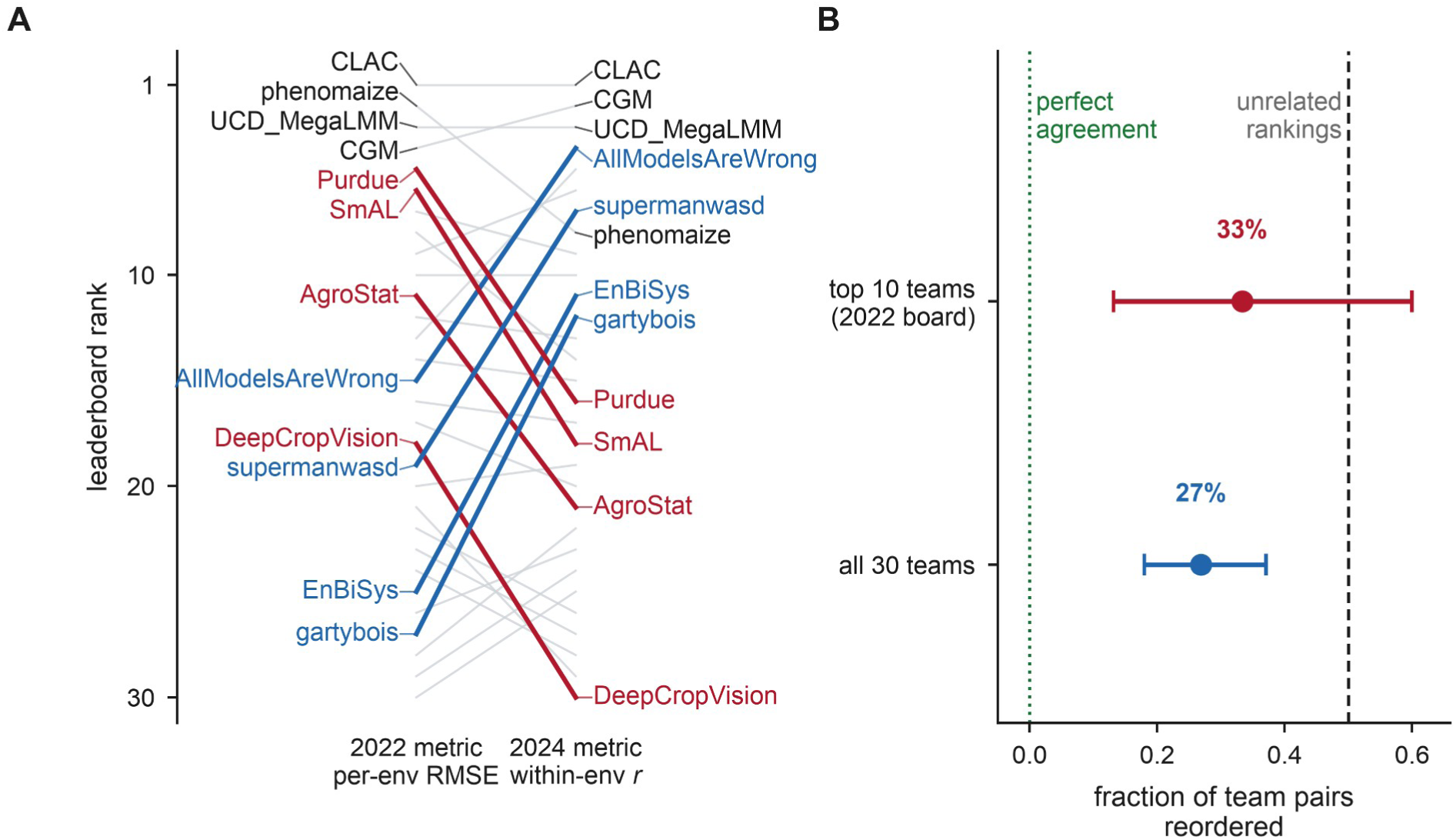
Changing the official metric reorders the G2F leaderboard. (A) Ranks of the 30 teams under the 2022 metric (mean per-environment RMSE) and the 2024 metric (mean within-environment Pearson *r*); the eight teams moving ten or more places are drawn in colour (red, falling; blue, rising). (B) Pairwise reversal rate with team-bootstrap 95 % intervals, for all 435 team pairs and for the 45 pairs among the top ten of the 2022 leaderboard. Dashed line, unrelated rankings; dotted line, perfect agreement.

The reversal is concentrated between metric classes: the two Tier I–II metrics reverse 9.4 % of team pairs against each other and the fourteen error-magnitude metrics a median of 5.1 % (0.0–9.4 %) among themselves, but the two groups a median of 27.6 % (26.9–29.0 %) against each other. Ranking the 30 teams by the Tier I metric alone differs from the 2022 official leaderboard by a median of 5 places (maximum 16) but from the 2024 one by 2 (maximum 8).

The same reversal appears in the constructed panels: 16–25 % of base-method pairs reverse, and 35–47 % once the miscalibrated variants are added (Table 1; Fig. 3B; Supplementary Table S4). The competition’s rate, 26.9 % across 30 independent submissions, lies between the two: the added miscalibration raises the rate but does not create it.

**Fig. 3.**
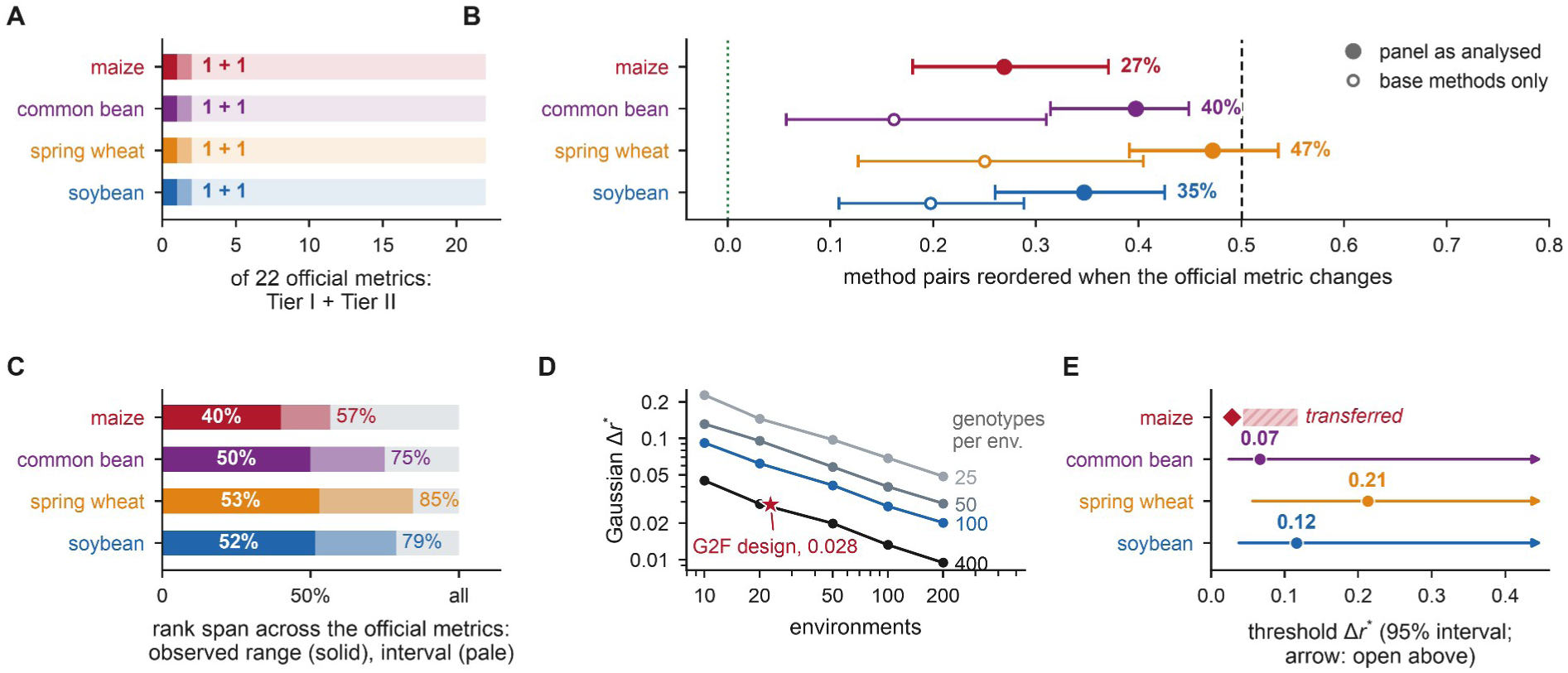
The answers across four datasets. (A) Official metrics in Tier I (dark) and Tier II (light). (B) Reversal rate between the two official metrics, with cluster-bootstrap 95 % intervals; filled, panel as analysed; open, base methods only. (C) Rank span across the official metrics: observed range (solid) and 95 % interval (pale). (D) Gaussian Δ*r*\* at 10 % selection against the number of environments, by genotypes per environment (Supplementary Table S11); star, the G2F design. (E) Threshold Δ*r*\* among base methods with two-way bootstrap 95 % intervals (arrow, upper limit beyond the observed range). Maize: the Gaussian value at the competition’s design (diamond, 0.028) and the range after scaling by the panels’ observed-to-Gaussian ratios (0.04–0.12).

A rank inherits this instability. Across the 22 official metrics a G2F team’s rank spans a median of 12 of 30 places (40 %), and the winning team’s spans first to seventh (Fig. 4); in the three panels the observed range spans 50–53 % of the board (Table 1; Fig. 3C; Supplementary Table S16). On the G2F board the 95 % rank interval is wider, a median of 17 places (57 %; Fig. 4), and its width comes from mixing metric classes rather than from the number of metrics: de-duplicating the 22 metrics to the six distinct quantities they represent leaves it at 17 places, whereas within the error-magnitude family, or within the two Tier I–II metrics, it narrows to 7 and 8 places (Supplementary Table S8).

**Fig. 4.**
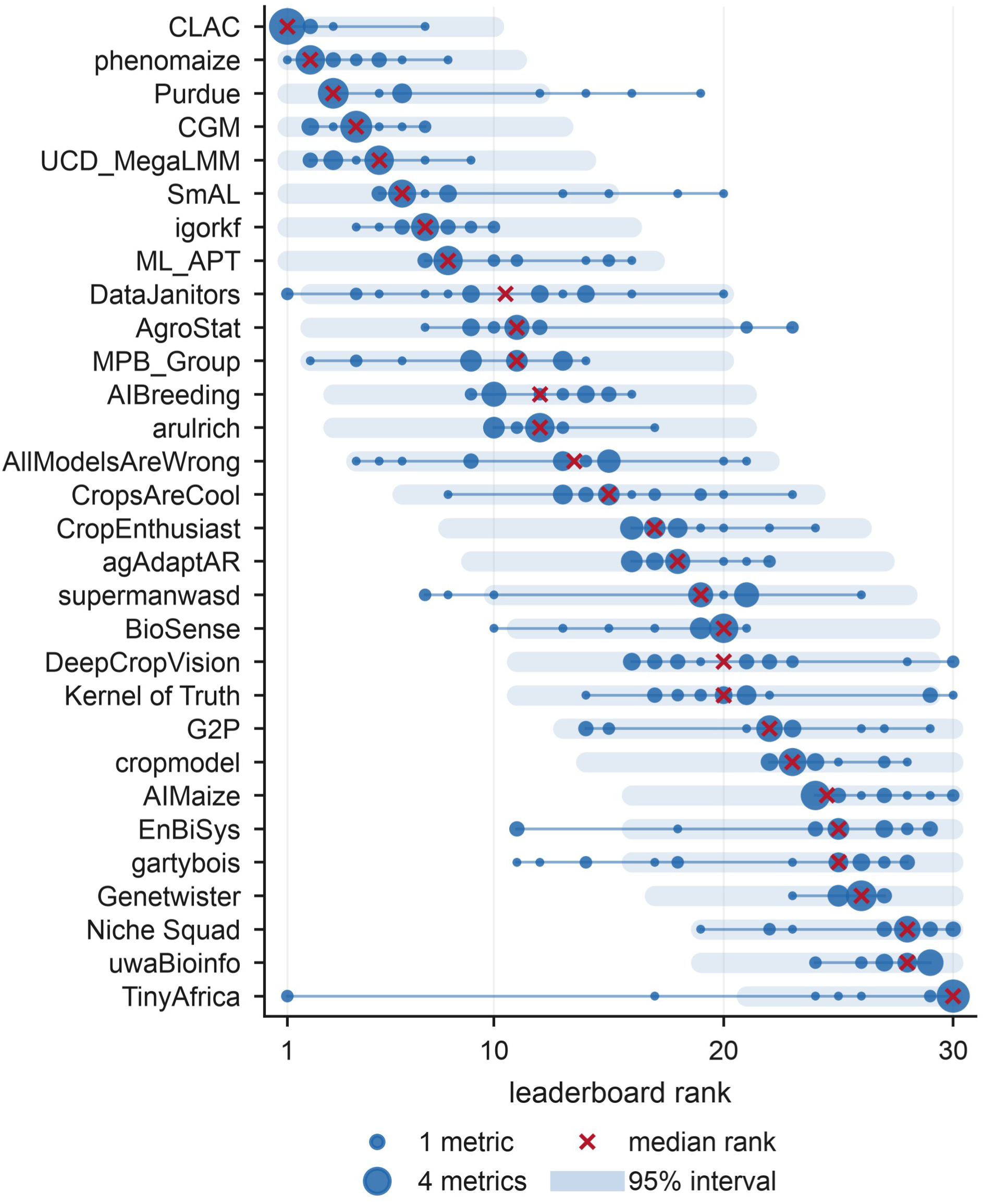
A published rank depends on the metric. For each of the 30 G2F teams, blob area is the number of the 22 official metrics placing the team at that rank; ×, median rank; pale band, 95 % interval. The median observed range is 12 of 30 places (interval, 17), and the first-placed team takes ranks 1 to 7.

### 3.3 How large an accuracy gain must be: larger than the gaps at the top of the leaderboard

Accuracy is used here as a surrogate for selection gain, which earns its place only if a change in it implies a change in the endpoint it stands for [64,65]; the smallest such effect is the surrogate threshold effect [66]. Our analogue is the gap Δ*r*\* in mean within-environment Pearson correlation, the scale on which the leaderboard is published, at which the more accurate of two methods also selects better material with probability 0.95. Δ*r*\* is thus the resolution of a trial: the smallest accuracy gap that a trial of a given design reliably converts into better observed selection.

At equal accuracy that probability is one half by symmetry, whatever the surrogate; what matters is how fast it rises (Fig. 5A). At 10 % selection the logistic estimate of Δ*r*\* is **0.066 in common bean, 0.213 in spring wheat and 0.116 in soybean** (isotonic check 0.063, 0.157 and 0.128; Fig. 3E). Two-way bootstrap lower limits are 0.024, 0.056 and 0.038; the upper limit lies beyond the observed range in all three panels, where 49 %, 12 % and 24 % of resamples do not reach 0.95. Estimating the threshold out of sample, with the accuracy gap taken from one half of the environments and the realised differential from the other, gives 0.126 in common bean, 0.253 in spring wheat and 0.179 in soybean — larger than the in-sample values in every panel, and conservative in the direction that matters here: a trial resolves less than the fitted threshold suggests (Supplementary Table S5).

**Fig. 5.**
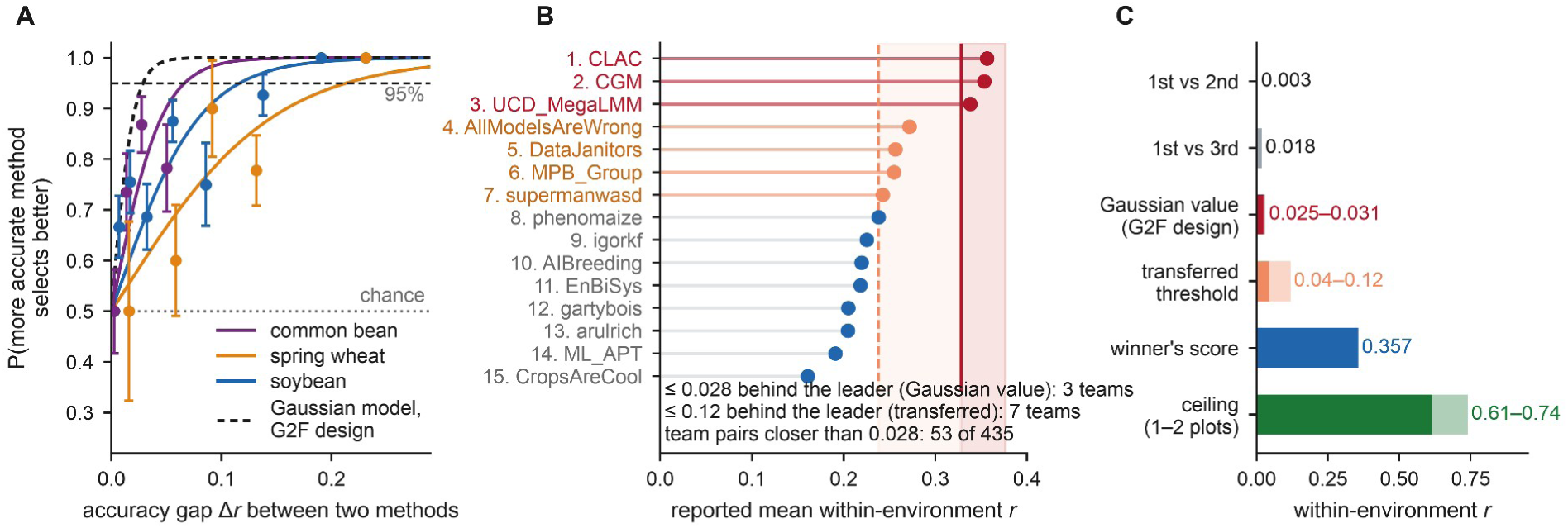
How large an accuracy gain must be before the more accurate method reliably selects better material. (A) Probability that the method with the higher mean within-environment Pearson *r* also has the higher realised selection differential at 10 % selection, against the accuracy gap, for base-method pairs in common bean (133 pairs), spring wheat (132) and soybean (324); points, binned proportions ± 1 SE; lines, logistic fits; dashed, the Gaussian model at the G2F design, which reaches 0.95 at 0.028. (B) Published mean_pearson_r of the top 15 G2F teams; red, within the Gaussian value (0.028) of the winner; orange, within the transferred upper value (0.12). (C) Leaderboard gaps, thresholds, the winner’s score and the ceiling — 0.61 if the scored value is a single plot and 0.74 if a two-plot mean, estimated from 114,234 plots of the competition’s training data — on one within-environment *r* scale.

For maize, whose five verified submissions give ten pairs, the threshold is obtained through the design. Gaussian panels matched to each constructed panel’s methods, genotypes per environment, environments and accuracy range give 0.043, 0.066 and 0.028, and the observed values are 1.5, 3.2 and 4.2 times these (Supplementary Table S10). At the competition’s design — 30 methods, 445 genotypes per environment, 23 environments, accuracies spanning the teams’ — the Gaussian value is 0.028 (0.025–0.031 as method errors go from independent to 80 % shared), and scaling by the panels’ ratios gives 0.04–0.12. The ratio rises with the number of environments across the three panels (1.5, 3.2 and 4.2 at 10, 109 and 169 environments) but is not ordered by genotypes per environment or by the number of genotype–environment cells, and three panels cannot separate the number of environments from the other ways in which they differ; we therefore report 1.5–4.2 as an empirical bracket observed on real panels rather than as a correction with a known cause.

#### Two statements hold under every estimate

First and second place, 0.003 apart in reported mean_pearson_r, are an order of magnitude below the Gaussian value and below every panel’s lower limit. And the first three places — CLAC 0.357, CGM 0.353, UCD_MegaLMM 0.338, spread 0.018 — lie within the Gaussian value of one another (Fig. 5B). Across the whole board, 53 of the 435 team pairs (12 %) lie closer than the Gaussian value and 229 (53 %) closer than the transferred upper value (Fig. 5B). In the Gaussian model, the design grid (Fig. 3D) shows what drives the value: the number of genotype–environment cells *N*, however divided between genotypes and environments. At *f* = 0.10 it falls as about 3/√*N* (log–log slope −0.54, *R*² = 0.99), from 0.13–0.15 at 500 cells to 0.03 at 8,000–10,000; the constant is 2.1 at *f* = 0.20 and 4.3 at *f* = 0.05 (Supplementary Table S11). A programme can read its own reference value off that grid, as a lower bound: the three panels here needed 1.5–4.2 times it.

Plot error caps any within-environment correlation at √*H*²(*n*), with *H*²(*n*) the heritability of a mean of *n* plots: 0.61 for one plot and 0.74 for two in the G2F training data. Because the replication of the 2022 answer key is undocumented, both are used: the winning team reached 48–58 % of the ceiling, and the first-to-third gap is 2.5–3.0 % of it (Fig. 5C; Supplementary Section S6).

A significant difference in accuracy is not the same as a reliable difference in selection. A Meng test on the pooled cells declares 56 %, 81 % and 92 % of base-method pairs different in common bean, spring wheat and soybean, but in only 80–83 % of those pairs does the more accurate method also select better, and 5–30 % of them reach the panel’s threshold (Supplementary Section S22).

### 3.4 Whether the decision’s metric selects better material: in soybean, where the trial separates the leading methods

Whether an admitted metric also selects better material is an empirical question, answered by the outcome test (Section 2.4). The outcome orders methods reliably in all three panels (split-half reliability 0.869 in common bean, 0.647 in spring wheat, 0.786 in soybean), and stays above the 0.5 criterion without the added variants (0.642, 0.683, 0.586).

For within-environment selection the two families diverge sharply in soybean, where the correlation rules pick the two GBLUP models (Supplementary Section S17): the correlation family recovers 98–99 % of the available selection gain against 13–34 % for the error-magnitude family, and the method ranked first by mean_pearson_r selected material 0.172 phenotypic SD better than that ranked first by mean_RMSE (Supplementary Table S7). With environments resampled that gap excludes zero for both correlation rules — mean_pearson_r +65 [+22, +90] percentage points against mean_RMSE, mean_spearman_r +64 [+19, +88] (Supplementary Table S15). The other two panels separate the families less: in spring wheat the correlation rules recover 52–73 % against 3–29 % for the error-magnitude rules, whereas in common bean all six rules recover 55–69 % and mean_RMSE (60 %) trails neither correlation rule (55 % and 57 %; Supplementary Table S7). The added variants do not drive these results: every rule except NDCG@10 % selects material of identical realised value with or without them, and what changes is chiefly the baseline, the average method. Nor do the GBLUP models: without them the correlation family still leads in soybean (78–80 % against 27–31 %; Supplementary Table S24).

The contrast between soybean and common bean is consistent with their resolution (Section 3.3). In soybean the correlation rules pick the two GBLUP models, the panel’s most accurate methods (mean within-environment *r* up to 0.36 against a panel median of 0.17), and 22 % of method pairs differ by more than the panel resolves; the 17 common-bean base methods span only 0.37–0.45, and 4 of their 133 pairs (3 %) do (Supplementary Tables S27 and S30). In common bean every rule therefore picks among methods the trial cannot separate, as RMSE does in the Chinese maize population, whose pick is one of three leaders the trial cannot separate (Section 3.6). Spring wheat, with 24 % of pairs resolvable, favours the correlation rules on point estimates only.

Repeated for a decision that reads calibration, the test could have shown that the criterion simply favours correlations; it does not. For that decision, mean_RMSE leads both correlation rules in common bean and spring wheat, by 8–113 percentage points, with the interval excluding zero in spring wheat and, at this sample size, not quite in common bean; in maize, from five submissions, mean_pearson_r leads instead (Supplementary Section S8). In spring wheat, on point estimates, the family that selects better therefore reverses with the decision. In soybean the order does not reverse: mean_RMSE (85 %) trails both mean_pearson_r (98 %) and mean_spearman_r (98 %) even under the absolute target (interval including zero), because the two GBLUP models pair the panel’s highest accuracy with near-best calibration (within-environment slopes 0.92 and 0.90, other base methods at most 0.63; mean RMSE within 1.5 % of the lowest), so the rules that identify them lead under both decisions. Admissibility is a validity requirement for the decision it is derived from, not a performance guarantee against every other decision.

Selecting on genotype means across environments is a different decision; its admissible metric, the rank correlation of predicted with observed genotype means, is none of the 22 official metrics. Repeated for this decision, the outcome test finds no detectable difference between mean_spearman_r and that metric where the outcome is reliable — −12 [−17, +8] percentage points in common bean and −1 [−13, +6] in soybean — and in soybean both lead mean_RMSE, by 40 and 42 percentage points, with intervals excluding zero (Supplementary Section S15).

Between the Tier I and Tier II rules the outcome test does not favour the admissible one: mean_spearman_r recovers within two percentage points of mean_pearson_r in common bean and soybean but 52 % against 73 % in spring wheat, and the top-weighted NDCG@10 % ranges from 33 % in spring wheat to 99 % in soybean. **Admissibility guarantees validity for the decision, not the best pick in every panel** (Section 3.6; Supplementary Table S24).

### 3.5 What replicates across species

Four species, two trait classes, competition submissions and constructed methods, and designs from 10 to 184 environments and 16 to 445 genotypes each give consistent answers (Table 1): the partition (Fig. 3A); reversal of 27–47 % of method pairs when the official metric changes, 16–27 % counting only unaugmented methods; a rank spanning 40–53 % of the board; and a surrogate threshold whose lower confidence limit, 0.024–0.056, exceeds the gap between the competition’s first two places.

Across 52 dataset × trait combinations from six datasets in four crops, the RMSE–Pearson reversal rate was not detectably associated within datasets with genomic heritability, the G×E share or the between-environment correlation of genotype values (Holm-adjusted *P* 0.40–0.42; Supplementary Section S21), so a trait’s genetic parameters give no grounds for skipping a check of whether the ranking of methods changes with the metric.

### 3.6 From criterion to verdict: a Chinese maize population

Applied to 17 methods predicting ear weight in the CUBIC population (1,400 recombinant inbred lines at five sites; 6,699 genotype–environment cells per method), GPverdict (Fig. 6A) recovers the same partition from the population’s own predictions. It ranks an ensemble of random forest, gradient boosting and a multilayer perceptron first by mean within-environment rank correlation (0.341), with random forest (0.340) and gradient boosting (0.323) inside the smallest gap the trial resolves, 0.036 (Fig. 6B). Splitting lines within sites, the method each rule picks recovers 79–90 % of the attainable selection gain (outcome reliability 0.80), the most for RMSE, whose pick, random forest, is one of the three tied leaders. The verdict is to choose among the three leaders on cost or simplicity (Supplementary Table S25). The same population scored on all 23 of its traits is shown in Fig. 6D and Supplementary Table S19.

**Fig. 6.**
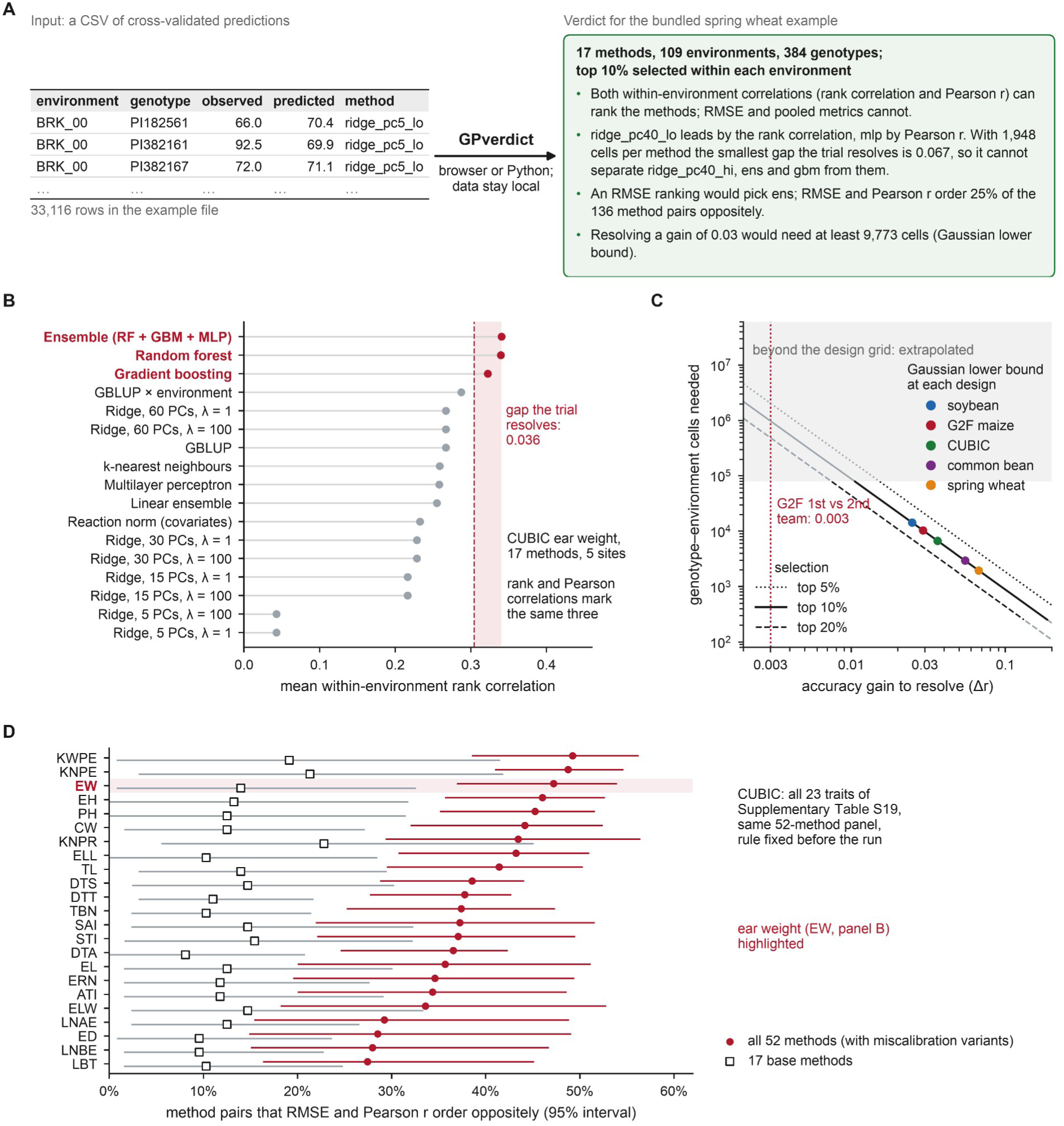
GPverdict turns the criterion into a verdict. (A) What GPverdict does: from a table of cross-validated predictions (one row per environment, genotype and method; the first rows of the tool’s bundled spring wheat example are shown) it returns a verdict, in the browser or in Python, without uploading data. The verdict shown is condensed from the tool’s report (version 1.1.1) on that example with the web page’s settings (top 10 % within each environment, at least 10 genotypes per environment); screenshots of the page and of the report’s verdict section are in Supplementary Fig. S5. (B) CUBIC ear weight: 17 methods ranked by mean within-environment rank correlation; shaded, the gap the trial resolves (0.036, Gaussian lower bound), within which three methods (red) cannot be separated. (C) Genotype–environment cells needed to resolve an accuracy gain at 5 %, 10 % and 20 % selection: black inside the Gaussian design grid (250–80,000 cells), grey extrapolated. Points, the Gaussian lower bound at each dataset’s design (observed thresholds are larger; Fig. 3E); dotted line, the gap between the first two G2F teams. (D) The same population scored on all 23 of its traits: share of method pairs that RMSE and Pearson *r* order oppositely, with 95 % intervals, for the 52-method panel (dots) and its 17 base methods (squares); ear weight, the trait of panel B, is highlighted (Supplementary Table S19).

## 4. Discussion

### 4.1 A criterion that adjudicates rather than describes

Disagreement between metrics is well documented; the central result here is that it can be adjudicated from the decision, and the partition of Section 3.1 follows before any data are examined.

The partition bears directly on current practice. Root mean squared error, mean absolute error, the coefficient of determination and the squared correlation all fail the criterion, as does the metric that determined the 2022 ranking. The organisers recommended Pearson correlation for breeding-focused competitions [16]; the family adopted for 2024 passes the affine test, and its rank-based member passes the full test. The failure is not a technicality: mean_r2_pearson, an official competition metric, cannot distinguish a method from its exact reverse, although the two differ by 1.6 phenotypic standard deviations in the material they advance (Section 3.1). Nor was the criterion built to prefer rank-based measures: when the decision becomes a fixed target value, which reads calibration, mean_RMSE overtakes both correlation rules in spring wheat, by 95 and 113 percentage points with intervals excluding zero, and leads them on point estimates in common bean (Section 3.4; Supplementary Table S15).

The criterion formalises and extends an observation made for hybrid rye, that losses and metrics pooled across environments favour the environment means [35]. The decomposition of Section 2.4 states it exactly: in the three constructed panels a median of 94–95 % of the attainable square of pooled Pearson correlation is carried by the predicted environment means (Supplementary Section S20). Eckhoff et al. prefer MSED because it also registers shrinkage, the right property for a decision that reads the size of differences; for truncation within an environment that information cannot be used, which places MSED in Tier III.

Inadmissible metrics keep their own roles: RMSE measures accuracy on the scale of the trait, which matters for any decision against an absolute target, and calibration itself is read from the slope and mean bias. The criterion identifies where such a metric should not be used — to rank methods for within-environment selection, which is what leaderboards are read as doing.

### 4.2 What a trial can resolve

The accuracy gain a trial needs before the more accurate method reliably selects better is a property of the trial and not of the methods; in the Gaussian model it is set mainly by the number of genotype–environment cells, so an evaluation can be sized to the gain it must detect (Fig. 6C), and the three real panels needed 1.5–4.2 times the Gaussian value (Section 3.3). A test that two correlations differ [59] asks whether the accuracies differ, the threshold whether the difference changes the material selected. Relatedness among genotypes [37] and effects shared by the environments of one year [36] make real cells less informative than the Gaussian grid assumes, so 3/√*N* is a lower bound and the calibrated range is the one to plan with. The outcome tests in soybean and common bean fit the same picture: the admissible metrics selected better material where the leading methods stood clear of the rest by more than the trial resolves, and made little difference where they did not (Section 3.4).

### 4.3 Scope and extensions

The criterion is derived for single-trait truncation selection within an environment; selection on an index or at a fixed fraction fixes its own admissible class by the same construction (Section 2.1), as selection on genotype means does (Section 3.4). For maize the threshold comes from the design; a direct estimate becomes possible if the competition’s organisers release the full set of submissions.

The constructed panels differ from the competition in a way this paper documents itself (Supplementary Section S3): their methods share one feature space, the same marker principal components, and vary only in the learner, whereas the 30 maize submissions came from independent teams. On the maize board the four within-environment discrimination metrics are the official metrics least predictable from the other 21; the panels do not show this separation, and restricting a panel to 20 well-connected environments restores it in spring wheat but not in soybean (Supplementary Section S3).

### 4.4 What should change

Three recommendations follow, in line with recent calls for application-oriented benchmarking [67]; GPverdict carries out all three on a programme’s own cross-validated predictions.

Competitions and studies should state which decision their score represents and verify that the score is invariant to its transformations and sensitive to their reversal. The test excludes twenty of the twenty-two metrics here.

Rankings should be published with the metric that produced them, the decision that metric represents and, as is standard when learning algorithms are compared [68], an interval; a reported improvement should come with the shortlist it would change, set against the design’s reference threshold.

For within-environment selection, methods should be ranked by the within-environment correlations — the rank correlation, which the criterion admits, and the Pearson correlation, which calibration cannot move and over which the outcome test did not favour the admissible one (Section 3.4) — not by an error-magnitude statistic. A correlation pooled across environments should not rank methods for within-environment selection: it moves by 0.85 under transformations that leave every such decision unchanged.

## 5. Conclusions

What a breeding programme needs from an accuracy ranking is not the ranking but the size of difference that can change the material it advances. For single-trait selection within environments, the decision itself fixes which metrics are admissible: those depending on the predictions only through their within-environment ranking, changing when it reverses. Of the twenty-two official metrics of a major maize competition, that is the mean within-environment rank correlation; the within-environment Pearson correlation is unmoved by calibration only, and the other twenty, including the metric that decided the 2022 ranking, fail the criterion. The choice is consequential for rankings: it reorders a quarter of team pairs. An accuracy gain must reach at least 0.03 in within-environment correlation at the competition’s design before the more accurate method reliably selects better, and 0.04–0.12 once calibrated to real panels, against 0.003 between the first two places. Resolution also helps to read the outcome tests: in soybean, where the leading methods stood clear of the rest by more than the trial resolves, the admissible correlations selected better material than error-magnitude rankings, and in common bean and a Chinese maize population, where they did not, every rule picked material of similar value.

### Practical guidance

For a breeding programme choosing a prediction model: (i) first check what the trial can resolve — at 10 % selection the gap is at least about 3/√*N* for *N* genotype–environment cells (4.3/√*N* at 5 % and 2.1/√*N* at 20 %), the real panels here needed 1.5–4.2 times that at 10 %, and the Gaussian lower bound is about 3,500 cells for a gain of 0.05 and 10,000 for 0.03; (ii) rank candidate models by the within-environment rank and Pearson correlations, or by the realised selection differential at the programme’s own selection fraction, and keep RMSE, calibration slope and bias as diagnostics; and (iii) treat models within the resolvable gap as indistinguishable, choose among them on cost or simplicity, and adopt a model upgrade only if the trial can resolve it. GPverdict performs these steps in a web browser.

## CRediT authorship contribution statement

**Guohao Lv:** Conceptualization, Methodology, Software, Validation, Formal analysis, Investigation, Data curation, Visualization, Writing – original draft, Writing – review & editing. **Renzhe Zheng:** Methodology, Software, Validation, Formal analysis, Investigation, Writing – review & editing. **Lichuan Gu:** Supervision, Project administration, Funding acquisition, Writing – review & editing.

## Declaration of generative AI and AI-assisted technologies in the writing process

During the preparation of this work the authors used Claude (Anthropic), under the authors’ direction, to check the numbers reported in the text against the analysis outputs and to translate the text into English and polish its language to improve readability. The authors take full responsibility for the content of the published article.

## Data and code availability

All data are public. Maize: the Genomes to Fields 2022 competition data (doi:10.25739/tq5e-ak26) and the official metrics of every submission [16]; the 2024 leaderboard is at EvalAI (challenge 2376) [63]. Common bean: doi:10.7910/DVN/XCD67U; spring wheat: doi:10.5061/dryad.wstqjq2z0; soybean: SoyBase (https://www.soybase.org) and Wartha et al. [50]. The sources and licences of every dataset, including the Chinese maize, DROPS (doi:10.15454/IASSTN) and SoyNAM data of Supplementary Sections S11 and S21, are listed in DATA_SOURCES.md of the code repository.

Scripts reproducing the analyses, with the derived data, are archived at Zenodo (https://doi.org/10.5281/zenodo.23051852) [69] and maintained at https://github.com/nblvguohao/gp-decision-criterion. GPverdict (version 1.1.1), with example data, is archived at Zenodo (https://doi.org/10.5281/zenodo.23051854) [60], maintained at https://github.com/nblvguohao/gpverdict and available as a web tool at https://nblvguohao.github.io/gpverdict/.

## Acknowledgements

We thank the Genomes to Fields Initiative and the organisers of the 2022 genotype-by-environment prediction competition for releasing the data and the complete table of official metrics, the teams who deposited their prediction files, and the groups who made the common bean, spring wheat and soybean data public.

