## Supplementary Material for "A decision-based criterion and a resolution threshold for ranking genomic prediction models within trials"

### Contents

| Item | Title |
|---|---|
| Section S1 | Problem fingerprint for selection under genotype-by-environment interaction |
| Table S1 | The fingerprint |
| Section S2 | Per-metric invariance, order sensitivity and leave-one-metric-out coverage |
| Table S2 | Metric-level values, maize and soybean |
| Section S3 | Which official metrics can be predicted from the others |
| Fig. S1 | Leave-one-metric-out coverage for each of the 22 official metrics |
| Fig. S2 | The two-dimensional metric taxonomy |
| Section S4 | Reversal rate between all pairs of official metrics |
| Fig. S3 | Pairwise reversal rate, 22 × 22 official metrics |
| Section S5 | Robustness checks |
| Table S3 | Noise null: disagreement against the sampling variability of a single metric |
| Table S4 | Reversal rates with and without the miscalibration variants |
| Table S5 | The surrogate threshold: estimates, checks and sensitivity |
| Table S6 | Rank-interval width: environment resampling against metric choice |
| Table S7 | Outcome test, all scoring rules and datasets |
| Table S8 | Rank-interval width under de-duplicated and family-restricted metric sets |
| Table S9 | The shortfall below $i_k \cdot r$: Gaussian-copula benchmark |
| Table S10 | Surrogate threshold: observed against its normal-theory counterpart |
| Table S11 | Design grid for the normal-theory threshold |

| | |
|---|---|
| Table S12 | Metrics from recent multi-environment studies under the invariance and order tests |
| Section S6 | The ceiling depends on the replication of the scored value |
| Table S13 | The ceiling on the within-environment correlation as a function of replication |
| Section S7 | Paired rank invariance: does a decision-neutral transformation reorder the submissions? |
| Table S14 | Probability that the ranking of the five verified submissions changes |
| Section S8 | Changing the decision |
| Table S15 | Outcome test under the within-environment and an absolute-threshold decision |
| Section S9 | The observed rank range across the official metrics |
| Table S16 | Observed rank range and rank interval, all four datasets |
| Section S10 | Tied predictions at the selection boundary |
| Table S17 | Frequency of exact ties and of ties straddling the top-10 % boundary |
| Section S11 | A fifth dataset: Chinese maize (CUBIC) |
| Table S18 | Principal results for CUBIC ear weight |
| Table S19 | The CUBIC population scored on all 23 agronomic traits |
| Section S12 | Composition of the constructed method panels |
| Table S20 | Base methods and settings in each panel |
| Section S13 | Which submission stands for a team |
| Table S21 | The leaderboard quantities under five selection rules |
| Section S14 | Tied predictions and the shortfall below $i_k \cdot r$ |
| Table S22 | The shortfall below $i_k \cdot r$ under three tie-breaks |
| Section S15 | Selection on genotype means across environments |
| Table S23 | Outcome test for selection on genotype means |
| Section S16 | Withdrawn panels and exploratory analyses |
| Section S17 | Panel composition and the outcome test |
| Table S24 | Outcome test under changes to panel composition |
| Section S18 | GPverdict: implementation, validation and the CUBIC verdict |
| Table S25 | GPverdict on CUBIC ear weight |
| Table S26 | GPverdict against the published values |
| Section S19 | Methods and results details for the main text |
| Section S20 | Pooled metrics decomposed into an environment-mean part and a within-environment part |
| Table S27 | Pooled Pearson correlation and pooled MSE decomposed |

| | |
|---|---|
| Section S21 | Genetic architecture and the disagreement between metrics |
| Table S28 | Genetic parameters of the 52 combinations |
| Table S29 | Tests of the pre-specified plan |
| Section S22 | The resolution threshold and tests that compare two correlations |
| Table S30 | Pairs declared different, and whether the more accurate method selects better |
| Table S31 | Smallest accuracy gap a pooled Meng test declares, against 3/√N |
| Section S23 | Population structure of the constructed panels |
| Table S32 | Marker structure of the three panels |
| Fig. S4 | Population structure of the three constructed panels |
| Fig. S5 | The GPverdict web page running its bundled example |

### Section S1. Problem fingerprint for selection under genotype-by-environment interaction

**Proof of the proposition of Section 2.1.** *A metric $M(\hat{y}, y)$ is invariant under all strictly increasing within-environment transformations of $\hat{y}$ if and only if it depends on $\hat{y}$ only through the within-environment rank pattern of $\hat{y}$, ties included.* Sufficiency is immediate, since such a transformation leaves that pattern unchanged. For necessity, take two prediction vectors sharing the same rank pattern in every environment. Each environment carries finitely many predictions, so the map sending the *j*-th smallest value of the first to the *j*-th smallest of the second is strictly increasing on the observed support and extends to a strictly increasing function on the line; applying it environment by environment carries the first vector to the second, and invariance forces the metric to take the same value on both. The transformation group is the product over environments of the strictly increasing maps of the line, $\prod_e G_e$, and the within-environment rank pattern is its maximal invariant; the statement is the standard characterisation of invariant functions through one. It concerns transformations of $\hat{y}$ at fixed $y$, and assumes the metric is defined on the vectors compared — a metric undefined for some of them, such as a within-environment regression slope when the predictions in an environment are constant, is outside its scope.

The fingerprint below was constructed following the problem-fingerprint approach of Maier-Hein et al. [S1]. Its purpose is to make explicit which properties of the decision constrain metric choice. The emboldened row is the constraint from which the admissibility criterion in Section 2.1 of the main text is derived; the others record context that a reader may wish to check against their own programme.

***Table S1. Problem fingerprint.***

| **Fingerprint dimension** | **Value for genomic selection under G×E** | **Consequence for metric choice** |
|---|---|---|
| Decision the prediction feeds | Advance the top *k* genotypes, *k* not fixed in advance; cull the rest | Not a regression problem; absolute predicted values are never read off |
| Unit the decision is made over | Genotype, **within one environment** | Metrics must aggregate within environment, never pool; between-environment variance is nuisance |
| Comparison set | Only genotypes grown in the same | A method that ranks environments well but |

| | | |
|---|---|---|
| | environment | genotypes poorly is worthless for selection |
| Selection intensity | Typically the top 5–20 % | Accuracy in the middle of the distribution carries almost no value |
| Value function | Expected selection differential $i \cdot r \cdot \sigma$ under joint normality of prediction and phenotype | Under that model only the within-environment correlation enters; bias and scale do not appear |
| **Required invariance** | **Any strictly increasing within-environment transform leaves the decision unchanged; reversing the ranking inverts it** | **A valid metric must be invariant to every strictly increasing within-environment transform and must change when the ranking is reversed — the criterion used throughout this work** |
| Cost asymmetry | Discarding an elite line ≫ advancing a mediocre one | Symmetric squared error weights errors in the middle of the distribution as heavily as errors among the selected |
| Data structure | Unbalanced genotype × environment table with missing cells | Pooling weights environments by plot count; the aggregation scheme is a substantive choice |
| Reference standard | Plot yield measured with error; within-environment repeatability $H^2 \approx 0.38$ per plot, 0.55 for a two-plot mean | The ceiling on any correlation metric is $\sqrt{H^2} \approx 0.61–0.74$ depending on the replication scored against, not 1 |

### Section S2. Per-metric invariance, order sensitivity and leave-one-metric-out coverage

Drift is the median relative change in a metric under a transformation of the predictions that cannot change any selection decision: a per-environment affine map (maize, five verified submissions, 200 replicates; soybean, the 26 base methods, 20 replicates) or a random strictly increasing map of the within-environment average ranks (maize, 120 replicates). A metric is order-sensitive if it changes by at least $10^{-3}$ when every environment's predictions are reversed. Tier I, monotone-invariant and order-sensitive; Tier II, affine-invariant and order-sensitive only; Tier III, all others. Coverage is the leave-one-metric-out coverage of the 95 % rank interval of Section 2.4 of the main text. Tables produced by analysis/crosscrop/code/partition.py and analysis/crosscrop/code/build_tableS2.py.

Three points are worth noting when reading Table S2. First, the Tier I and II metrics drift by at most $2 \times 10^{-16}$ under the affine test, while the smallest affine drift among the order-sensitive Tier III metrics is 0.10 in the maize test of Table S2 and 0.075–0.148 across all five datasets in the 20-replicate partition test, so the classification does not depend on the exact tolerance. Second, mean_r2_pearson is unmoved by the affine test in every dataset but, like pooled $r^2$, unchanged by reversal, so both are Tier III; in spring wheat mean_r2_pearson gives the best base method and the same method with its predictions reversed within every environment the same score (Fig. 1D of the main text). Third, the tiers are identical, metric by metric, in maize, common bean, spring wheat, soybean and Chinese maize (partition_metrics.csv), as they must be, since they follow from the definitions.

***Table S2. Metric-level values.***

| **Metric** | **Aggregation** | **Quantity** | **Affine drift,** | **Monotone drift,** | **Order-** | **Tier** | **Affine drift,** | **Coverage,** | **Coverage,** |
|---|---|---|---|---|---|---|---|---|---|

| | | | **maize** | **maize** | **sensitive** | | **soybean** | **maize** | **soybean** |
|---|---|---|---|---|---|---|---|---|---|
| mean_spearman_r | within-env | discrimination | 0.0 | 0.0 | yes | **I** | 0.0 | 0.70 | 0.86 |
| mean_pearson_r | within-env | discrimination | $2.0 \times 10^{-16}$ | $1.7 \times 10^{-1}$ | yes | **II** | 0.0 | 0.70 | 0.88 |
| mean_r2_pearson | within-env | discrimination | $2.5 \times 10^{-16}$ | $3.0 \times 10^{-1}$ | **no** | III | 0.0 | 0.80 | 0.85 |
| mean_lineRegressSlope | within-env | discrimination | $1.0 \times 10^{-1}$ | $3.5 \times 10^{-1}$ | yes | III | $7.5 \times 10^{-2}$ | 0.67 | 0.89 |
| mean_relative_RMSE | within-env | error magnitude | $7.5 \times 10^{-1}$ | 1.70 | yes | III | 1.16 | 1.00 | 1.00 |
| mean_RMSE | within-env | error magnitude | $7.9 \times 10^{-1}$ | 2.15 | yes | III | 1.22 | 1.00 | 1.00 |
| mean_normalized_RMSE | within-env | error magnitude | $7.8 \times 10^{-1}$ | 2.13 | yes | III | 1.22 | 1.00 | 1.00 |
| mean_realative_MAE | within-env | error magnitude | $8.7 \times 10^{-1}$ | 1.98 | yes | III | 1.33 | 1.00 | 1.00 |
| RMSE | pooled | error magnitude | $9.3 \times 10^{-1}$ | 2.05 | yes | III | 1.51 | 1.00 | 1.00 |
| normalized_RMSE | pooled | error magnitude | $9.3 \times 10^{-1}$ | 2.05 | yes | III | 1.51 | 1.00 | 1.00 |
| relative_RMSE | pooled | error magnitude | $9.3 \times 10^{-1}$ | 2.05 | yes | III | 1.51 | 1.00 | 1.00 |
| mean_normalized_MAE | within-env | error magnitude | $9.2 \times 10^{-1}$ | 2.53 | yes | III | 1.39 | 1.00 | 1.00 |
| MAE | pooled | error magnitude | $9.5 \times 10^{-1}$ | 2.72 | yes | III | 1.56 | 1.00 | 1.00 |
| normalized_MAE | pooled | error magnitude | $9.5 \times 10^{-1}$ | 2.72 | yes | III | 1.56 | 1.00 | 1.00 |
| realative_MAE | pooled | error magnitude | $9.5 \times 10^{-1}$ | 2.72 | yes | III | 1.56 | 1.00 | 1.00 |
| mean_MAE | within-env | error magnitude | $9.3 \times 10^{-1}$ | 2.58 | yes | III | 1.41 | 1.00 | 1.00 |
| r2_pearson | pooled | discrimination | $9.1 \times 10^{-1}$ | $9.3 \times 10^{-1}$ | **no** | III | $8.4 \times 10^{-1}$ | 0.97 | 0.85 |
| lineRegressSlope | pooled | discrimination | $9.7 \times 10^{-1}$ | 1.26 | yes | III | $8.7 \times 10^{-1}$ | 0.93 | 0.79 |
| pearson_r | pooled | discrimination | $9.7 \times 10^{-1}$ | $8.4 \times 10^{-1}$ | yes | III | $6.1 \times 10^{-1}$ | 0.97 | 0.85 |

| | | | | | | | | | |
|---|---|---|---|---|---|---|---|---|---|
| | | tion | | | | | | | |
| spearman_r | pooled | discrimination | 1.02 | $8.5 \times 10^{-1}$ | yes | III | $6.3 \times 10^{-1}$ | 1.00 | 0.83 |
| mean_r2_score | within-env | error magnitude | 3.78 | 10.99 | yes | III | 5.37 | 1.00 | 1.00 |
| r2_score | pooled | error magnitude | 20.65 | 60.66 | yes | III | 30.49 | 1.00 | 1.00 |

The metric names are reproduced verbatim from the competition's published table, including the spellings realative_MAE and realative_RMSE.

### Section S3. Which official metrics can be predicted from the others

Holding out each metric in turn and asking how well the remaining 21 predict a team's rank under it, the four lowest coverages in the maize data are mean_lineRegressSlope 0.67, mean_spearman_r 0.70, mean_pearson_r 0.70 and mean_r2_pearson 0.80. The fifth lowest is 0.93, and nothing lands in the 0.133-wide band between them (Fig. S1). All four of the bottom group are computed within environments and none is an error-magnitude quantity; all fourteen error-magnitude metrics are recovered with coverage 1.00.

Both axes of the taxonomy are needed to see this, and each cell behaves as the mechanism predicts (Fig. S2). mean_r2_score is computed within environments yet is perfectly predictable, because $R^2$_score = 1 − SSE/SST is an error-magnitude quantity in a within-environment coat. Pooled Pearson *r* is likewise predictable, because pooled correlation is dominated by between-environment variance — 58.0 % of the total in the 2022 maize test set (between_env_share.py). Only within-environment discrimination is a genuinely separate axis.

**Sensitivity to classification.** We state the result as an ordering rather than as a property of a metric class, because the class-level version depends on how two metrics are assigned. Grouping the calibration slope with the discrimination metrics, as we do, gives a clean class boundary; removing it does not, because the within-environment slope is itself the least predictable metric and moving it into the comparison group lowers that group's minimum to 0.67. Classifying $R^2$_score as discrimination likewise destroys the boundary, although that assignment contradicts its algebraic form. The empirical ordering is unaffected by either choice.

**Across datasets.** The separation holds in maize, where the four within-environment discrimination metrics are the least predictable, well clear of the rest (0.67–0.80 against at least 0.93 for every other metric). It does not hold in any of the three constructed panels: the within-environment discrimination range meets or sits inside the range of every other metric (common bean 0.79–0.87 and spring wheat 0.81–0.87, against 0.87–1.00; soybean 0.85–0.89 against 0.79–1.00). The constructed panels differ from the maize competition in one respect that may contribute: their methods share one feature space, the same marker principal components, and vary only in the learner applied to it, whereas the 30 maize submissions came from independent teams using different modelling strategies. Restricting a panel to well-connected environments gives a mixed answer: a subset of 20 soybean environments with a mean pairwise Jaccard overlap of the genotype sets of 0.396 (0.066 in the full series) does not restore separation, whereas a subset of 20 spring-wheat environments (0.382 against 0.152) does, with the within-environment discrimination metrics at 0.77–0.85 against at least 0.87 for every other metric (analysis/crosscrop/code/connectivity_test.py, connectivity.csv). The between-environment share of variance — 58.0 % in maize, 67.8 % in common bean, 50.6 % in spring wheat and 69.5 % in soybean — does not order the datasets by whether they separate (analysis/crosscrop/code/between_env_share.py). The main-text conclusions rest on the invariance criterion rather than on this analysis.

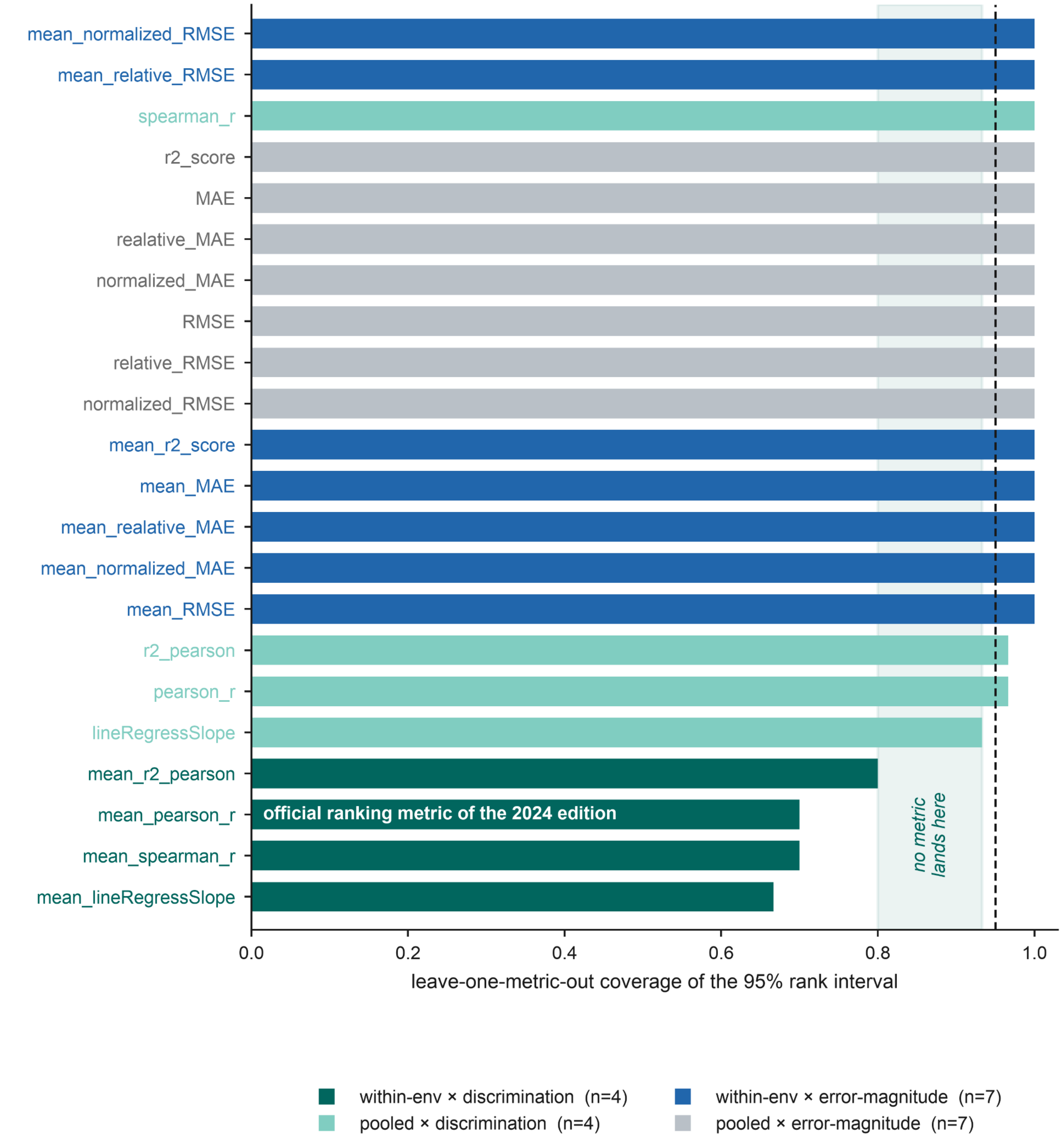

**Fig. S1.** Leave-one-metric-out coverage of the 95 % rank interval for each of the 22 official metrics, coloured by aggregation and quantity. Dashed line, nominal 95 %. Shaded band, the empty interval separating the four lowest metrics from the remaining eighteen — a descriptive gap, not a test. White label, the metric adopted as the official ranking score for the 2024 edition.

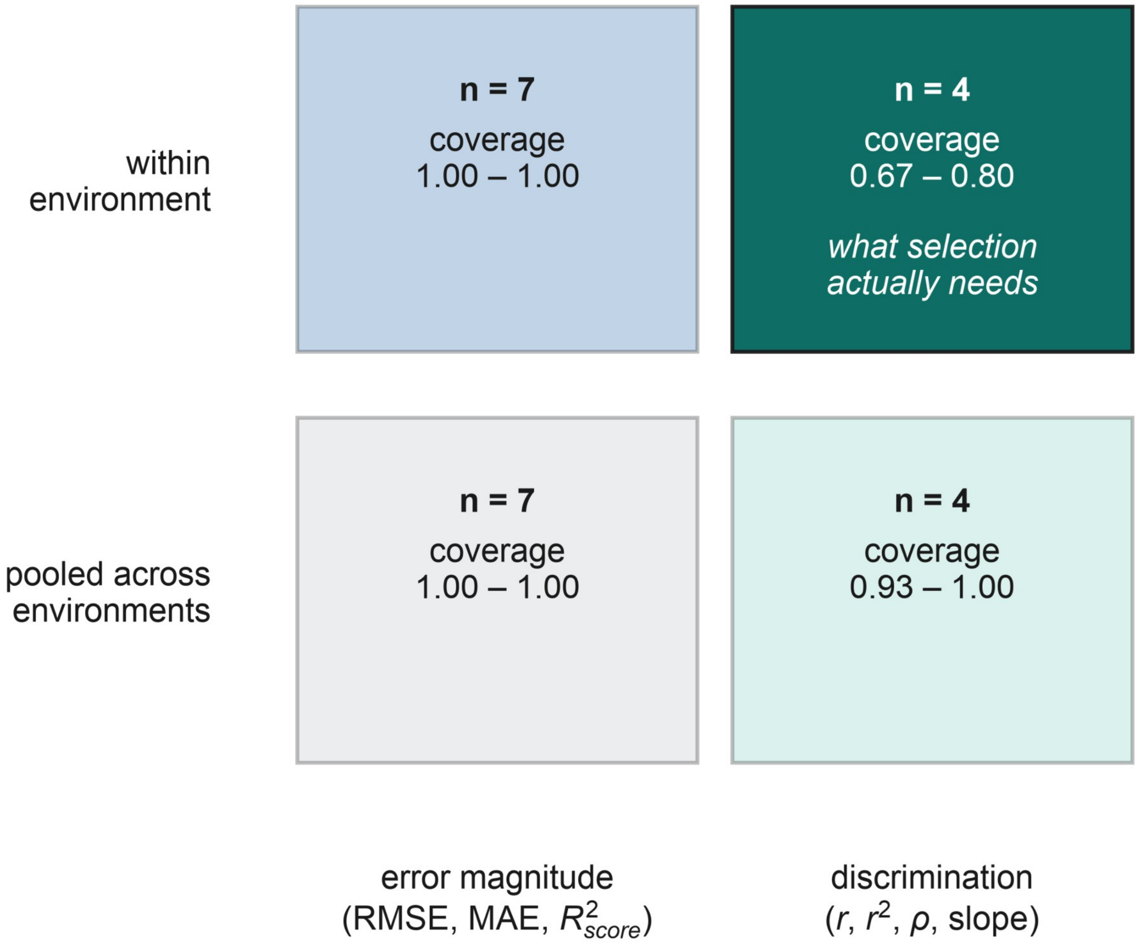

**Fig. S2.** The 22 official metrics cross-classified by aggregation (pooled versus within-environment) and quantity (error magnitude versus discrimination), with the range of leave-one-metric-out coverage in each cell. Only the within-environment discrimination cell behaves differently, and it is the cell selection decisions depend on.

### Section S4. Reversal rate between all pairs of official metrics

Across all 231 pairs of the 22 official metrics the median reversal rate is 11.7 %; 56.7 % of metric pairs reverse more than 10 % of team pairs and 39.4 % reverse more than 20 %. Excluding the two calibration-slope metrics, whose orientation involves a judgement call, leaves 190 pairs with a median of 9.2 % and 26.8 % above 20 %.

Each pair was tested against the null that it reverses at most 5 % of team pairs. Team pairs share teams and are not independent trials, so the standard error comes from resampling teams (4,000 replicates, self-pairs excluded) and the test is a one-sided Wald $z$. Ten pairs never reverse — the RMSE variants among themselves, the MAE variants among themselves, and pooled Pearson $r$ against pooled $r^2$ — so their bootstrap standard error is zero; they are the strongest possible evidence *for* the null and are scored $P = 1$. After Benjamini–Hochberg control across the 231 pairs, 129 remain significant at FDR 5 % and 84 at FDR 1 %; a percentile version of the same bootstrap gives 131 and 117. For the two official metrics the adjusted $P$ is $2.8 \times 10^{-5}$ (percentile, $6 \times 10^{-4}$). The 231 tests are positively dependent, which BH tolerates, but they are not 231 independent findings. Script: analysis/g2f_leaderboard/code/reversal_tests.py.

Ordering the metrics by hierarchical clustering of the reversal matrix (Fig. S3) recovers the same structure as Section S3: the error-magnitude metrics form a mutually consistent block, and the within-environment discrimination metrics do not join it.

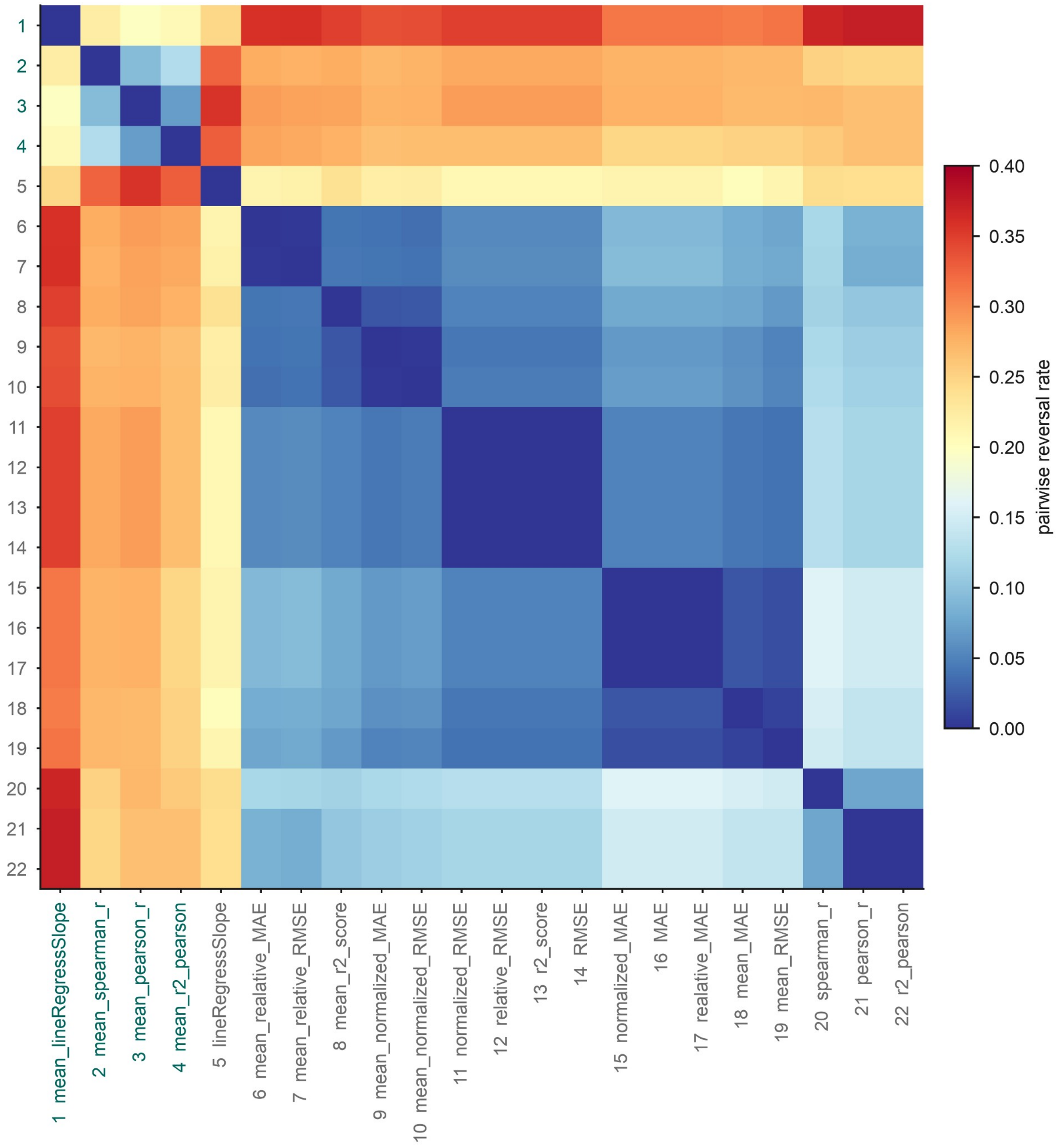

**Fig. S3.** Pairwise reversal rate between every pair of the 22 official metrics, ordered by hierarchical clustering. Within-environment discrimination metrics are labelled in teal. The axes are numbered: 1 mean_lineRegressSlope; 2 mean_spearman_r; 3 mean_pearson_r; 4 mean_r2_pearson; 5 lineRegressSlope; 6 mean_realative_MAE; 7 mean_relative_RMSE; 8 mean_r2_score; 9 mean_normalized_MAE; 10 mean_normalized_RMSE; 11 normalized_RMSE; 12 relative_RMSE; 13 r2_score; 14 RMSE; 15 normalized_MAE; 16 MAE; 17 realative_MAE; 18 mean_MAE; 19 mean_RMSE; 20 spearman_r; 21 pearson_r; 22 r2_pearson.

### Section S5. Robustness checks

***Table S3. Noise null.***

The disagreement between two metrics is compared against the sampling variability of a single metric, estimated by splitting environments into two disjoint halves (20 splits) and computing the reversal rate between the same metric evaluated on each half. Because each half carries only half the environments, this estimate of the noise floor is conservative. Script: analysis/crosscrop/code/robustness.py.

| Dataset | Signal: reversal between the two official metrics | Noise: mean_RMSE, half vs half | Noise: mean_pearson_r, half vs half | Signal / mean noise |
|---|---|---|---|---|
| spring wheat | 0.472 | 0.029 | 0.107 | **7.0×** |
| soybean | 0.347 | 0.131 | 0.060 | 3.6× |
| common bean | 0.397 | 0.090 | 0.170 | 3.0× |

***Table S4. Reversal rates with and without the miscalibration variants.***

Panels fitted by least squares are homogeneous in calibration in a way real competition entries are not, and were augmented as described in Section 2.3 of the main text. Removing the augmentation leaves 17–26 base methods per dataset. Pairs tied on either metric are excluded; intervals resample methods (with the augmentation removed, each method family is a single method), 2,000 replicates. Script: analysis/crosscrop/code/unaugmented_panels.py.

| Dataset | Methods, augmented / base | Reversal, augmented | Reversal, base | 95 % CI, base |
|---|---|---|---|---|
| spring wheat | 52 / 17 | 0.472 | 0.250 | 0.127–0.405 |
| soybean | 66 / 26 | 0.347 | 0.198 | 0.108–0.288 |
| common bean | 52 / 17 | 0.397 | 0.162 | 0.057–0.310 |

The augmentation raises the rate in all three panels but does not create it. The maize value from 30 independent submissions, 0.269, lies between the unaugmented (0.162–0.250) and augmented (0.347–0.472) ranges.

***Table S5. The surrogate threshold: estimates, checks and sensitivity.***

$\Delta r^*$ is the accuracy gap at which the logistic fit through one-half reaches 0.95 (Section 2.4 of the main text); "not reached", beyond the largest observed gap. Base methods are the primary analysis; "all" adds the miscalibration variants, with tied pairs dropped. The 95 % interval resamples method families and environments together (500 replicates); its upper limit is not identified when more than 2.5 % of resamples do not reach 0.95, and the share of such resamples is given. Out-of-sample: $\Delta r$ from one random half of the environments and $\Delta g$ from the other, 200 splits pooled. P(agree | $\Delta r < 0.01$): the share of such pairs in which the more accurate method has the larger realised differential, with the number of pairs in brackets; overlap: the mean top-10 % shortlist overlap of the pairs with $|\Delta r| < 0.01$, a set at least as large (39, 226, 11, 95, 60 and 180 pairs by row) because it is taken before the outcome-tie filter. Scripts: analysis/crosscrop/code/threshold_model.py, threshold_estimate.py.

| **Dataset** | **Methods** | **Pairs** | ***f* = 0.05** | ***f* = 0.10** | ***f* = 0.20** | **Isotonic, *f* = 0.10** | **95 % CI, *f* = 0.10** | **Resamples not reaching 0.95** | **Out-of-sample** | **P(agree** | **Δ*r* < 0.01)** | **Overlap** |
|---|---|---|---|---|---|---|---|---|---|---|---|---|
| common bean | base (17) | 133 | not reached | **0.066** | 0.037 | 0.063 | 0.024 – not identified | 49 % | 0.126 | 0.50 (36) | 0.64 | |
| common bean | all (52) | 1,278 | 0.193 | 0.056 | 0.046 | 0.061 | 0.031–0.201 | 0 % | 0.134 | 0.55 (220) | 0.56 | |
| spring wheat | base (17) | 132 | 0.208 | **0.213** | 0.116 | 0.157 | 0.056 – not identified | 12 % | 0.253 | 0.43 (7) | 0.72 | |
| spring wheat | all (52) | 1,277 | 0.200 | 0.208 | 0.130 | 0.166 | 0.081 – not identified | 11 % | 0.291 | 0.44 (88) | 0.40 | |
| soybean | base (26) | 324 | not reached | **0.116** | 0.042 | 0.128 | 0.038 – not identified | 24 % | 0.179 | 0.67 (60) | 0.76 | |
| soybean | all (66) | 2,096 | 0.223 | 0.094 | 0.049 | 0.082 | 0.038–0.223 | 1 % | 0.128 | 0.66 (180) | 0.60 | |

At $\Delta r = 0$ the agreement probability is one half by symmetry for any surrogate, so the equal-accuracy column tests nothing about accuracy; its spread around 0.5 reflects the few pairs available. The threshold falls as selection relaxes from $f = 0.10$ to $f = 0.20$ in all six rows, because a larger shortlist is more forgiving of small accuracy differences; its upper limit is identified in two of the six rows. We therefore do not propose a universal value; the transferable quantity is the design-based reference of Tables S10 and S11.

***Table S6. Rank-interval width: environment resampling against metric choice.***

Environment resampling: environments drawn with replacement, keeping their multiplicity, and methods ranked by mean_pearson_r alone (400 replicates); median 95 % percentile-interval width, from analysis/crosscrop/code/robustness.py. Metric choice: the interval of Section 2.4 across the 22 official metrics, on all environments, as in Table S16 (rank_census.py). The two are different constructions and do not add; their ratio shows which source of uncertainty is larger.

| **Dataset** | **Environments** | **Environment resampling, metric fixed** | **Metric choice, environments fixed** | **Methods** |
|---|---|---|---|---|
| soybean | 184 | 12 (18 %) | 52 (79 %) | 66 |

| | | | | |
|---|---|---|---|---|
| spring wheat | 109 | 12 (23 %) | 44 (85 %) | 52 |
| common bean | 10 | 19 (37 %) | 39 (75 %) | 52 |

In every panel the choice of metric moves a method's rank 2.1 to 4.3 times as far as resampling environments does. Collecting more environments narrows the sampling component and leaves the metric component untouched.

***Table S7. Outcome test.***

Realised selection gain recovered by each scoring rule's first choice (the mean over methods tied for first), as the fraction of the distance from an average method to the best method on the held-out half (top 10 % of genotypes, 400 environment splits). Because that best method is chosen on the held-out half itself, the fractions compare rules on a common scale rather than measure absolute efficiency. The reliability row is the split-half reliability of the realised selection differential; datasets below 0.5 are not used for comparison. Script: analysis/crosscrop/code/does_it_help.py.

| **Scoring rule** | **Tier** | **common bean** | **soybean** | **spring wheat** | **common bean, base** | **soybean, base** | **spring wheat, base** | **maize (5 submissions)** |
|---|---|---|---|---|---|---|---|---|
| *outcome reliability* | | *0.869* | *0.786* | *0.647* | *0.642* | *0.586* | *0.683* | — |
| mean_pearson_r | II | 55 % | 99 % | 73 % | 11 % | 99 % | 78 % | 13 % |
| mean_spearman_r | I | 57 % | 98 % | 52 % | 14 % | 97 % | 59 % | 71 % |
| mean_NDCG@10% | I | 69 % | 99 % | 33 % | 39 % | 99 % | 55 % | 67 % |
| mean_RMSE | III | 60 % | 34 % | 29 % | 20 % | 19 % | 38 % | 7 % |
| mean_MAE | III | 57 % | 23 % | 24 % | 15 % | 5 % | 34 % | 4 % |
| mean_r2_score | III | 68 % | 13 % | 3 % | 37 % | −6 % | 15 % | −39 % |

Five of the six rules select material of identical realised value with and without the added variants in all three panels; mean_NDCG@10% does so in soybean but not in common bean or spring wheat, and the best method on the held-out half is unchanged except in spring wheat (0.394 against 0.388 SD). The base columns' percentages therefore move mainly because the baseline, the average method, differs. In realised units, the first choice of mean_pearson_r exceeds that of mean_RMSE by 0.172 phenotypic SD in soybean (0.540 against 0.368). In maize, where the rule must choose among five submissions from three teams, the outcome's reliability cannot be estimated.

***Table S8. Rank-interval width under de-duplicated and family-restricted metric sets.***

Fourteen of the 22 official metrics are variants within the error-magnitude family, so the width reported in the main text could in principle be a property of the metric panel rather than of the leaderboard. The rank intervals of Section 3.2 were therefore recomputed on de-duplicated and family-restricted metric sets. The winner's interval is that of CLAC, first under the 2022 official metric. Produced by analysis/g2f_leaderboard/code/interval_sensitivity.py.

| **Metric set** | **Metrics** | **Median 95 % interval width** | **Fraction of the 30-team board** | **Winner's interval** |
|---|---|---|---|---|
| All 22 official metrics (main text) | 22 | 17 | 57 % | [1, 10] |
| Six distinct quantities, within environment | 6 | 17 | 57 % | [1, 10] |
| Six distinct quantities, pooled | 6 | 15 | 50 % | [1, 9] |
| Family-balanced: 2 error, 2 correlation, 1 slope | 5 | 19.5 | 65 % | [1, 11] |
| Error-magnitude family only | 14 | 7 | 23 % | [1, 4] |
| Correlation family only | 6 | 15 | 50 % | [1, 9] |
| Tier I and II metrics only | 2 | 8 | 27 % | [1, 4] |

Removing the redundancy leaves the width unchanged: six distinct quantities computed within environment give the same median width of 17 places as all 22 metrics, and balancing the families widens it. The width contracts only when the panel is restricted to a single class, to 7 places within the error-magnitude family and 8 for the two Tier I–II metrics. This is the between-class structure of Section 3.2 arriving by a second route, and it rules out metric multiplicity as the source of the reported width.

***Table S9. The shortfall below $i_k \cdot r$: Gaussian-copula benchmark.***

Standardised selection differential of each verified G2F submission's own top-*k* pick, against two benchmarks. *Normal benchmark*: $i\_\mathrm{fin}(n,k) \cdot r$, normal margins and the observed Pearson correlation, at the environment's actual *n* and *k*. *Copula benchmark*: a Gaussian copula carrying the environment's observed yield distribution and its observed Spearman correlation, so that both the margin and the rank dependence are held at their observed values. The SE is that of the gap against the copula benchmark, over the 23 environments (21 for Niche Squad); the SEs of the gap against $i_k \cdot r$ quoted in Section S19 are 0.019–0.046. A normal-margin control isolating the effect of the distribution's shape from that of matching on rank differs from the copula benchmark by 0.003–0.009 SD, so the shape of the yield distribution accounts for almost none of the difference between the two benchmarks; the submission for which they diverge most, EnBiSys 243568, is the one whose Pearson correlation most overstates its rank agreement. The five submissions come from three teams (Section 2.2 of the main text). Produced by analysis/crosscrop/code/copula_bridge.py.

| Submission | $f$ | Observed | Normal benchmark | Copula benchmark | Gap vs normal | Gap vs copula | SE, gap vs copula |
|---|---|---|---|---|---|---|---|
| KernelOfTruth 244906 | 0.05 | 0.031 | 0.292 | 0.260 | -0.261 | -0.229 | 0.041 |
| KernelOfTruth 244906 | 0.10 | 0.155 | 0.249 | 0.223 | -0.094 | -0.068 | 0.028 |
| KernelOfTruth 244906 | 0.20 | 0.112 | 0.199 | 0.179 | -0.087 | -0.067 | 0.019 |
| KernelOfTruth 244944 | 0.05 | -0.154 | 0.201 | 0.215 | -0.355 | -0.368 | 0.045 |
| KernelOfTruth 244944 | 0.10 | -0.068 | 0.171 | 0.184 | -0.239 | -0.251 | 0.031 |
| KernelOfTruth 244944 | 0.20 | 0.012 | 0.137 | 0.147 | -0.124 | -0.135 | 0.020 |
| KernelOfTruth 244953 | 0.05 | 0.129 | 0.268 | 0.258 | -0.139 | -0.130 | 0.044 |
| KernelOfTruth 244953 | 0.10 | 0.119 | 0.228 | 0.221 | -0.109 | -0.102 | 0.024 |
| KernelOfTruth 244953 | 0.20 | 0.129 | 0.182 | 0.177 | -0.053 | -0.048 | 0.020 |
| EnBiSys 243568 | 0.05 | -0.029 | 0.448 | 0.165 | -0.477 | -0.194 | 0.036 |
| EnBiSys 243568 | 0.10 | 0.073 | 0.382 | 0.142 | -0.308 | -0.068 | 0.025 |
| EnBiSys 243568 | 0.20 | 0.038 | 0.305 | 0.114 | -0.267 | -0.076 | 0.015 |
| NicheSquad 244985 | 0.05 | -0.062 | 0.240 | 0.162 | -0.301 | -0.223 | 0.043 |
| NicheSquad 244985 | 0.10 | 0.033 | 0.204 | 0.138 | -0.171 | -0.105 | 0.035 |
| NicheSquad 244985 | 0.20 | 0.054 | 0.163 | 0.111 | -0.110 | -0.057 | 0.017 |
| all five, $f$ = 0.05 | | -0.017 | 0.290 | 0.212 | -0.307 | -0.229 | |
| all five, $f$ = 0.10 | | 0.063 | 0.247 | 0.181 | -0.184 | -0.119 | |
| all five, $f$ = 0.20 | | 0.069 | 0.197 | 0.146 | -0.128 | -0.077 | |

Every gap remains negative under the copula benchmark, at 4.1–5.4 standard errors on average, so between three fifths and three quarters of the shortfall is accounted for by neither the finite population, the shape of the yield distribution, nor the choice of correlation coefficient. With the margin and the rank correlation both held at their observed values, what remains can only be a departure of the dependence itself from the Gaussian form.

***Table S10. Surrogate threshold: observed against its normal-theory counterpart.***

The threshold estimator of Section 2.4 applied to Gaussian panels matched to each dataset's design — the number of base methods, the median genotypes per environment and environments per method entering the analysis, and accuracies spread over the panel's own 2.5th–97.5th percentile range (accuracies below 0.01 set to 0.01 in the simulation) — at $f = 0.10$. $\lambda$ is the fraction of error variance the methods share; $\lambda = 0$ makes them err independently. Each value is the median of 40 simulated panels. For maize the configuration is the competition's: 30 teams, their accuracy range, 445 genotypes and 23 environments. Produced by analysis/crosscrop/code/threshold_theory.py.

| **Dataset** | **Methods** | **Genotypes per environment** | **Environments** | **Accuracy range** | **Observed** | **$\lambda = 0$** | **$\lambda = 0.5$** | **$\lambda = 0.8$** | **Observed / theory ($\lambda = 0.5$)** |
|---|---|---|---|---|---|---|---|---|---|
| common bean | 17 | 344 | 10 | 0.37 to 0.45 | 0.066 | 0.048 | 0.043 | 0.036 | 1.5× |
| spring wheat | 17 | 16 | 109 | −0.05 to 0.35 | 0.213 | 0.064 | 0.066 | 0.060 | 3.2× |
| soybean | 26 | 61 | 169 | 0.15 to 0.35 | 0.116 | 0.032 | 0.028 | 0.025 | 4.2× |
| maize (G2F design) | 30 | 445 | 23 | −0.04 to 0.35 | — | 0.031 | 0.028 | 0.025 | transferred: 0.044–0.118 |

The observed threshold exceeds its normal-theory counterpart in all three panels — 1.5× in common bean, 3.2× in spring wheat and 4.2× in soybean — and changing $\lambda$ moves the theoretical value little. The Gaussian model is therefore a lower reference for how much accuracy separation a design can resolve, which every real panel exceeded. The maize value is transferred by multiplying the competition's Gaussian value ($\lambda = 0.5$) by the smallest and largest of the three observed/theory ratios.

***Table S11. Design grid for the normal-theory threshold.***

Threshold $\Delta r^*$ at $f = 0.10$ under the Gaussian model, for 40 methods with accuracies spread over 0.05–0.45 and $\lambda = 0.5$; the median of 25 simulated panels. Values for $f = 0.05$ and $f = 0.20$ are in analysis/crosscrop/results/threshold_design.csv.

| **Genotypes per environment** | **10 env** | **20 env** | **50 env** | **100 env** | **200 env** |
|---|---|---|---|---|---|
| 25 | 0.228 | 0.145 | 0.098 | 0.069 | 0.049 |
| 50 | 0.132 | 0.096 | 0.058 | 0.040 | 0.029 |
| 100 | 0.092 | 0.062 | 0.041 | 0.028 | 0.020 |
| 400 | 0.045 | 0.029 | 0.020 | 0.013 | 0.009 |

The reference value falls with genotypes per environment, with environments and with relaxed selection, and the three trade off: 50 genotypes in 200 environments, 100 in 100 and 400 in 20 resolve about the same accuracy difference (0.028–0.029). The whole grid is summarised by $\Delta r^* \approx k/\sqrt{N}$, with $N$ the number of genotype–environment cells: $k = 4.3$, 3.0 and 2.1 at $f = 0.05$, 0.10 and 0.20 (log–log slopes −0.55, −0.54 and −0.54; $R^2 \geq 0.99$), and the formula comes within 18 % of every value in the grid. At $f = 0.10$, resolving 0.05 therefore takes about 3,500 cells and resolving 0.03 about 10,000, the size of the G2F 2022 test set (23 environments of about 445 genotypes). A programme can read its own reference value here; by Table S10, the three panels needed 1.5–4.2 times theirs.

***Table S12. Metrics from recent multi-environment studies under the invariance and order tests.***

Median relative change over 40 replicates per method and transform (one reversal); tolerance $10^{-3}$; Tier I at a fixed fraction for the set-based metrics.

| **Metric** | **Test** | **maize (5 verified submissions)** | **common bean** | **spring wheat** | **soybean** |
|---|---|---|---|---|---|
| rooted MSED | affine | $6.3\times10^{-3}$ | $4.3\times10^{-2}$ | $6.0\times10^{-2}$ | $4.9\times10^{-2}$ |
| | monotone | $1.6\times10^{-3}$ | $9.1\times10^{-2}$ | $5.2\times10^{-2}$ | $5.4\times10^{-2}$ |
| | reversal | $5.0\times10^{-2}$ | $3.9\times10^{-1}$ | $1.5\times10^{-1}$ | $1.6\times10^{-1}$ |
| | tier | III | III | III | III |
| MSEPD | affine | $1.0\times10^{-2}$ | $8.9\times10^{-2}$ | $1.2\times10^{-1}$ | $7.8\times10^{-2}$ |
| | monotone | $4.0\times10^{-3}$ | $1.7\times10^{-1}$ | $8.1\times10^{-2}$ | $8.1\times10^{-2}$ |
| | reversal | $9.6\times10^{-2}$ | $8.4\times10^{-1}$ | $5.4\times10^{-1}$ | $4.5\times10^{-1}$ |
| | tier | III | III | III | III |
| coincidence index, 10 % | affine | 0 | 0 | 0 | 0 |
| | monotone | 0 | 0 | 0 | 0 |
| | reversal | 4.8 | 1.4 | 1.8 | 1.6 |
| | tier | I | I | I | I |
| coincidence index, 20 % | affine | 0 | 0 | 0 | 0 |
| | monotone | 0 | 0 | 0 | 0 |
| | reversal | 5.1 | 1.7 | 1.5 | 1.8 |
| | tier | I | I | I | I |
| coincidence index (denominator as in Fernandes et al.), 10 % | affine | 0 | 0 | 0 | 0 |

| | | | | | |
|---|---|---|---|---|---|
| | monotone | 0 | 0 | 0 | 0 |
| | reversal | 4.8 | 1.4 | 2.1 | 1.6 |
| | tier | I | I | I | I |
| coincidence index (denominator as in Fernandes et al.), 20 % | affine | 0 | 0 | 0 | 0 |
| | monotone | 0 | 0 | 0 | 0 |
| | reversal | 5.1 | 1.7 | 1.7 | 1.8 |
| | tier | I | I | I | I |
| NDCG, 10 % | affine | 0 | 0 | 0 | 0 |
| | monotone | 0 | 0 | 0 | 0 |
| | reversal | $1.0\times10^{-1}$ | $3.0\times10^{-1}$ | $1.2\times10^{-1}$ | $1.1\times10^{-1}$ |
| | tier | I | I | I | I |
| NDCG, 20 % | affine | 0 | 0 | 0 | 0 |
| | monotone | 0 | 0 | 0 | 0 |
| | reversal | $8.2\times10^{-2}$ | $2.7\times10^{-1}$ | $9.7\times10^{-2}$ | $8.8\times10^{-2}$ |
| | tier | I | I | I | I |
| within-environment Spearman ρ | affine | 0 | $2.0\times10^{-6}$ | 0 | 0 |
| | monotone | 0 | 0 | 0 | 0 |
| | reversal | 2.0 | 2.0 | 2.0 | 2.0 |
| | tier | I | I | I | I |
| within-environment Pearson r | affine | $2.1\times10^{-16}$ | 0 | $1.9\times10^{-16}$ | $1.4\times10^{-16}$ |
| | monotone | $1.6\times10^{-1}$ | $7.5\times10^{-2}$ | $1.3\times10^{-1}$ | $9.3\times10^{-2}$ |
| | reversal | 2.0 | 2.0 | 2.0 | 2.0 |
| | tier | II | II | II | II |
| per-environment RMSE | affine | $9.1\times10^{-1}$ | 1.4 | 1.7 | 1.3 |
| | monotone | 2.2 | 3.6 | 4.4 | 3.7 |
| | reversal | 5.8 | 8.1 | 9.8 | 8.3 |
| | tier | III | III | III | III |

### Section S6. The ceiling depends on the replication of the scored value

A prediction can correlate with an observed value only as far as that value correlates with the genotypic mean it stands for. If the scored value averages $n$ plots, the bound is $\sqrt{H^2(n)}$ with $H^2(n) = \sigma^2_g/(\sigma^2_g + \sigma^2_e/n)$, so the ceiling describes the measurement, not the trait: as $n$ grows it approaches 1. We estimated $\sigma^2_g$ and $\sigma^2_e$ per environment from the G2F 2014–2021 training data (128,167 plots) by unbalanced one-way random-effects ANOVA [S2,S3], retaining the 192 environments with at least 20 entries, at least 20 error degrees of freedom and a positive genetic component — 114,234 plots (Table S13). Replicate effects are not removed, because plots of one hybrid occupy different replicates by design and that variation is part of what separates a plot from the genotypic mean; removing them raises the ceiling by 0.02 at $n = 1$.

Restricting the estimate to cells with two or more surviving plots gives $\sqrt{H^2} = 0.796$. That subset is 43.5 % of the 84,527 environment-by-hybrid cells and is enriched for checks and for cells without plot loss; estimating over all cells gives 0.740 at the same $n = 2$. The competition's answer key holds one value per environment-by-hybrid cell with no replicate count, and neither the competition data note nor the competition paper states how many plots stand behind it. A variance diagnostic — comparing each scored environment's within-environment variance with the training-year components at the same location — implies $n = 1.45$ (95 % CI 0.94–1.88 over the 23 environments), consistent with the same mixture of one- and two-plot cells as the training data (harmonic mean 1.29 plots per cell, 56.5 % single plots). It assumes the 2022 variance components match the 2014–2021 medians at each location and is biased upward by the quality filtering applied to the released file, so we report the ceiling as the interval implied by $n = 1$ and $n = 2$, 0.61 to 0.74, rather than as a point.

Common bean is subject to the same alignment question. Its published BLUEs rest on 2.97 plots per line per trial, so the reliability route's 0.773 describes a three-plot mean; re-estimating from the raw plots at each trial's own replication gives 0.794, and at a single plot 0.603 against maize's 0.614. The two species agree at matched replication, which is the check the bean data were brought in to provide.

***Table S13. The ceiling on the within-environment correlation as a function of the replication of the scored value.***

Median over 192 G2F environments (2014–2021 training data, 114,234 of its 128,167 plots) and 9 VEF common-bean trials; 95 % intervals from a bootstrap over environments (respectively trials), 2,000 draws. Shares use the competition's published mean_pearson_r: winner 0.3566 and first-to-third gap 0.0183. Script: analysis/crosscrop/code/heritability_ceiling_A8.py.

| **Plots behind the scored value $n$** | **Maize $H^2(n)$** | **Maize ceiling $\sqrt{H^2(n)}$** | **Bean ceiling** | **Winner's share of ceiling** | **1st-to-3rd gap as share** |
|---|---|---|---|---|---|
| 1 (single plot) | 0.377 | 0.614 [0.580, 0.645] | 0.603 [0.512, 0.728] | 58.0 % | 2.97 % |
| 1.29 (training, harmonic) | 0.439 | 0.663 [0.629, 0.692] | 0.651 | 53.8 % | 2.76 % |
| 1.45 (implied by the 2022 variance diagnostic) | 0.467 | 0.684 [0.650, 0.712] | — | 52.2 % | 2.67 % |
| 2 (two-plot mean) | 0.548 | 0.740 [0.709, | 0.730 [0.644, | 48.2 % | 2.47 % |

| | | | | | |
|---|---|---|---|---|---|
| | | 0.767] | 0.832] | | |
| 3 | 0.645 | 0.803 [0.777, 0.826] | 0.794 [0.718, 0.878] | 44.4 % | 2.27 % |
| 4 | 0.708 | 0.841 [0.818, 0.861] | 0.834 [0.766, 0.905] | 42.4 % | 2.17 % |
| → ∞ (genotypic mean) | 1 | 1 | 1 | 35.7 % | 1.83 % |
| *replicated cells only* | *0.634* | *0.796* | *0.773 (n ≈ 3)* | *44.8 %* | *2.30 %* |

### Section S7. Paired rank invariance: does a decision-neutral transformation reorder the submissions?

The invariance test of Section 2.4 perturbs one submission and measures how far each metric's value moves; the claims from Section 3.2 onwards concern the order of methods. The paired version applies an independent decision-neutral transformation to each of the five verified G2F submissions and asks whether the resulting ranking of the five differs from the untransformed ranking. The transformations are applied within environment and are strictly increasing there, so every submission's own top-*k* pick in every environment is unchanged by construction and any movement of the ranking is movement no selection decision can justify. Monotone transformations use average ranks, so that exactly tied predictions map to the same value; with ordinal ranks, which break ties, mean_spearman_r still never reorders the submissions, and for the other metrics the probability of a change moves by at most 0.01 and the pair-reversal rate by up to 0.19. 400 replicates; ten submission pairs. Script: analysis/g2f_leaderboard/code/paired_invariance.py.

***Table S14. Probability that the ranking of the five verified submissions changes.***

| **Metric** | **Tier** | **Affine: P(ranking changes)** | **Affine: pairs reversed** | **Monotone: P(ranking changes)** | **Monotone: pairs reversed** |
|---|---|---|---|---|---|
| mean_spearman_r | I | 0.000 | 0.000 | 0.000 | 0.000 |
| mean_pearson_r | II | 0.000 | 0.000 | 0.998 | 0.344 |
| mean_r2_pearson | III (order-insensitive) | 0.000 | 0.000 | 0.995 | 0.412 |
| the other 19 | III | 0.960–1.000 | 0.292–0.699 | 0.988–1.000 | 0.347–0.696 |

A metric that passes the value-level test never reorders the submissions under the corresponding class, and every metric that fails it reorders them in at least 96 % of replicates. The partition of Section 3.1 is therefore a partition of the rankings the metrics produce, not only of their values. mean_r2_pearson passes the affine test but is Tier III because it cannot distinguish a ranking from its reverse, which this test, applying only order-preserving maps, does not probe. The Monte-Carlo upper 95 % bound on the zero entries is 0.009.

### Section S8. Changing the decision

The outcome test of Section 3.4 judges scoring rules by the realised selection differential of a within-environment top-*k* pick, which is itself a within-environment rank functional. A reader may therefore ask whether the correlation family was bound to win. The test was rerun with the scoring rules, panels and split-half protocol unchanged and the decision replaced by one the criterion does not protect: advance every genotype–environment cell whose predicted value clears a fixed target, the 90th percentile of observed values in that dataset, and book the realised surplus over the target. A within-environment monotone transformation changes which cells clear the target, so this decision lies outside the scope conditions of Section 2.1. Ties on a rule's score are resolved by averaging over the tied methods. Intervals resample environments and split the distinct resampled environments between the two halves, so no environment contributes to both; the paired contrast is the difference of the two rules' split-averaged recoveries. Script: analysis/crosscrop/code/decision_swap.py.

***Table S15. Outcome test under the within-environment and an absolute-threshold decision.***

Recovery as in Table S7, from an independent set of splits; the last two columns are the paired difference in recovery between each correlation rule and mean_RMSE, in percentage points, so that the admissible metric (Tier I, mean_spearman_r) is contrasted and not only its family. Maize rows are shown for completeness: five submissions from three teams give no reliability estimate, and of their four paired contrasts only the Tier II one under the absolute threshold excludes zero.

| Dataset | Decision | Outcome reliability | mean_RMSE | mean_pearson_r | mean_spearman_r | Pearson − RMSE [95 % CI] | Spearman − RMSE [95 % CI] |
|---|---|---|---|---|---|---|---|
| common bean | absolute threshold | 0.88 | 97 % | 89 % | 70 % | −8 [−97, +1] | −27 [−118, +1] |
| spring wheat | absolute threshold | 0.93 | 99 % | −14 % | 4 % | −113 [−121, −97] | −95 [−130, −45] |
| soybean | within-environment top 10 % | 0.79 | 35 % | 99 % | 99 % | +65 [+22, +90] | +64 [+19, +88] |
| soybean | absolute threshold | 0.78 | 85 % | 98 % | 98 % | +13 [−6, +34] | +12 [−6, +33] |
| maize (5 submissions) | within-environment top 10 % | — | 6 % | 10 % | 74 % | +4 [−73, +119] | +68 [−29, +161] |
| maize (5 submissions) | absolute threshold | — | −3 % | 100 % | −25 % | +103 [+34, +125] | −22 [−91, +0] |

Under the within-environment decision both correlation rules lead mean_RMSE in soybean, with both intervals excluding zero. Under the absolute-threshold decision the correlation rules fall behind RMSE in common bean and spring wheat — decisively in spring wheat, where both intervals exclude zero, and on the point estimates only in common bean, where both intervals reach +1; in soybean they lead RMSE numerically even under this decision, though the interval includes zero. The outcome test therefore does not favour the correlation family by construction: in spring wheat the family that leads reverses with the decision, and in soybean neither the within-environment nor the absolute-threshold decision favours RMSE, because the two GBLUP models pair the panel's best discrimination with near-best calibration — within-environment slopes of 0.92 and 0.90 against at most 0.63 for any other base method, and a mean RMSE within 1.5 % of the lowest, which belongs to a *k*-nearest-neighbour model with far weaker discrimination (Section 3.4 of the main text). In maize the Tier II rule goes the other way under the absolute threshold (+103 [+34, +125]) while the Tier I rule does not (−22 [−91, 0]); with five methods the rule's first choice is one of five, and no reliability can be estimated. With a pooled top-10 % decision across environments, the error-magnitude family leads in spring wheat (mean_RMSE 94 % against 72 % and 78 % for the two correlation rules) and is closely matched by the correlation family in common bean; in soybean the correlation family leads even under this pooled decision (point estimates in decision_swap_d3.csv; a pooled quota cannot be resampled by environment).

### Section S9. The observed rank range across the official metrics

The rank interval of Section 3.2 pools its deviations over all method–metric cells, so its half-width $q$ is a single number and every method receives the same unclipped width $2q + 1$; methods differ only where the interval is clipped at rank 1 or rank $M$. Because it is fitted and evaluated on the same cells, the share of cells escaping it (the escape rate) is close to 5 % by construction, and its coverage would also require the metrics to be exchangeable. Table S16 therefore reports, beside it, the observed range of each method's ranks across the 22 official metrics, which needs neither assumption. Scripts: analysis/crosscrop/code/rank_census.py (the table) and analysis/g2f_leaderboard/code/ties_and_coverage.py (the leave-one-family-out coverages quoted below).

***Table S16. Observed rank range and rank interval, all four datasets.***

| Dataset | Methods | Median observed range (places) | Median observed interquartile span | Median interval width | Escape rate | Leader, observed | Leader, interval |
|---|---|---|---|---|---|---|---|
| maize | 30 | 12 (40 %) | 2 | 17 (57 %) | 4.2 % | [1, 7] | [1, 10] |
| common bean | 52 | 26 (50 %) | 9.0 | 39 (75 %) | 4.7 % | [1, 17] | [1, 26] |
| spring wheat | 52 | 27.5 (53 %) | 9.0 | 44 (85 %) | 4.6 % | [1, 18] | [1, 34] |
| soybean | 66 | 34 (52 %) | 11.0 | 52 (79 %) | 4.6 % | [1, 20] | [1, 36] |

The leader is the base method ranked first by mean_pearson_r; a miscalibration variant ties with its parent on that metric and is not counted as the leader. The small interquartile span beside the large range is the between-class structure of Section 3.2: most metrics place a method within a few ranks of one another, and the range comes from the metrics of the other family. The same structure breaks the exchangeability that coverage would assume. In maize, an interval calibrated on the 14 error-magnitude metrics ($q = 3$) covers the ranks the six correlation metrics give in 56 % of cases, and one calibrated on the 20 non-slope metrics covers the two slope metrics' ranks in 80 %; leaving out a single metric gives 94.2 %. The interval's centre, a method's median rank across metrics, is a rank-based aggregate of the kind that can favour misspecified models when used for model selection under skewed scores [S4]; it is used here only to describe how far a rank moves, and the mean within-environment rank correlation recommended in the main text is a per-environment score averaged across environments, not such an aggregate.

### Section S10. Tied predictions at the selection boundary

A strictly increasing transformation cannot separate exactly tied predictions, and the proposition of Section 2.1 accordingly characterises admissible metrics through the within-environment rank pattern with ties included. Where a tie straddles the *k*-th place the selected set is itself not unique. Table S17 counts how often that happens. Script: analysis/g2f_leaderboard/code/ties_and_coverage.py.

***Table S17. Frequency of exact ties and of ties straddling the top-10 % boundary.***

A case is one method in one environment.

| Dataset | Cases | Cells in an exact tie | Cases with any tie | Cases with a tie straddling the boundary |
|---|---|---|---|---|
| maize (5 verified submissions) | 115 | 3.7 % | 89.6 % | 4.3 % |
| common bean | 520 | 1.7 % | 51.7 % | 1.5 % |
| spring wheat | 5,668 | 0.3 % | 2.3 % | 0.1 % |
| soybean | 11,154 | 3.2 % | 13.6 % | 1.0 % |

### Section S11. A fifth dataset: Chinese maize (CUBIC)

CUBIC is a population of 1,404 recombinant inbred lines from 24 Chinese elite maize founders, grown at five Chinese sites in one season [S5]; per-site ear weight is from the published supplementary data [S6] and genotypes from the CropGS-Hub release of the same population [S7]. A 52-method panel was built on the same design as the common-bean and spring-wheat panels — five-fold cross-validation on genotypes, the same learners, GBLUP and the marker × environment GBLUP, and the same variants — with the differences listed in Table S20 (principal components 5–60 from thinned markers, a reaction norm on environmental covariates, and no marker-only ridge). Scripts: analysis/crosscrop/code/panel_china.py (panel) and summary_china.py (Table S18); provenance and licences: analysis/crosscrop/results/china_DATA_SOURCE.md. The phenotype table carries a CC BY-NC-ND licence, so neither it nor the derived panel is redistributed in the archive. Ear weight is analysed as the yield trait of the population. All 23 traits were put through the same 52-method panel; the driver is analysis/crosscrop/code/china_trait_sweep.py and the summary table is analysis/crosscrop/results/china_trait_sweep.csv. Reversal rates between the two official metrics run 27–49 % with a median of 37 %, every one with a 95 % interval above zero; counted on the 17 base methods alone they run 8–23 % with a median of 13 %, and the interval excludes zero in 19 of the 23. Ear weight gives 47 %, the third highest. No trait and no site was dropped: the smallest site carries 1,266 genotypes, far above the 25-genotype floor. The reversal is therefore not specific to the trait chosen, and its size for ear weight lies at the upper end of the traits' range.

***Table S18. Principal results for CUBIC ear weight.***

| Result | CUBIC maize |
|---|---|
| Environments / genotypes per environment | 5 / 1,336 |
| Official metrics in Tier I / II / III | 1 / 1 / 20 |
| Largest affine drift of a Tier I–II metric | 0.0 |
| Smallest affine drift of an order-sensitive Tier III metric | 0.15 |
| Reversal between the two official metrics [95 % CI] | 47.2 % [36.9, 53.9] |
| The same, unaugmented panel (17 methods) [95 % CI] | 14.0 % [0.8, 32.6] |
| Median rank-interval width | 43 of 52 (83 %) |
| Outcome reliability | 0.893 |
| Threshold $\Delta r^*$ | not estimable: five environments, eight shared required |

The partition is the same as in the four main datasets. Section 3.6 of the main text and Section S18 give GPverdict's verdict for the 17 base methods. The reversal rate equals the highest of theirs (range 27–47 %), and without the added variants lies slightly below the unaugmented panels and maize (16–27 %).

The trait sweep. Four rules were fixed before the sweep was run and are recorded, with a UTC timestamp, in analysis/crosscrop/results/china_trait_sweep_RUN_STARTED.txt: (R1) every trait of the phenotype table is included, none added and none removed; (R2) the 25-genotype environment floor of the main text applies, a site below it is dropped, and a trait left with fewer than two environments is dropped; (R3) a trait passes the quality gate if its best base method reaches a mean within-environment Pearson correlation of 0.15, a value set between the 0.08 and 0.11 of the two withdrawn panels and the 0.34 of ear weight; (R4) the table reports every trait attempted, whether or not it passes. The gate is reported for completeness and did not operate here: the lowest value across the 23 traits is 0.307. The sweep therefore does not test the gate's behaviour; it records that no CUBIC trait approaches the near-chance regime of Section S16. The 23 traits are one population in one season at five sites and are not independent replications of one another; what they establish is that the results above do not depend on which trait of this population is scored.

***Table S19. The CUBIC population scored on all 23 agronomic traits.***

| Trait | Best base r | Gate | Reversal (95 % CI) | Reversal, base (95 % CI) | Interval width | Reliability |
|---|---|---|---|---|---|---|
| ATI | 0.389 | pass | 0.343 [0.200, 0.486] | 0.118 [0.016, 0.291] | 68 % | 0.903 |
| CW | 0.410 | pass | 0.442 [0.320, 0.524] | 0.125 [0.016, 0.271] | 80 % | 0.886 |
| DTA | 0.468 | pass | 0.365 [0.246, 0.423] | 0.081 [0.000, 0.208] | 69 % | 0.925 |
| DTS | 0.468 | pass | 0.386 [0.288, 0.441] | 0.147 [0.024, 0.302] | 68 % | 0.959 |
| DTT | 0.479 | pass | 0.378 [0.277, | 0.110 [0.031, | 70 % | 0.926 |

| | | | | | | |
|---|---|---|---|---|---|---|
| | | | 0.427] | 0.217] | | |
| ED | 0.359 | pass | 0.285 [0.149, 0.491] | 0.096 [0.008, 0.236] | 64 % | 0.922 |
| EH | 0.482 | pass | 0.460 [0.356, 0.527] | 0.132 [0.000, 0.318] | 81 % | 0.879 |
| EL | 0.399 | pass | 0.357 [0.200, 0.511] | 0.125 [0.016, 0.301] | 72 % | 0.922 |
| ELL | 0.520 | pass | 0.432 [0.307, 0.510] | 0.103 [0.000, 0.285] | 78 % | 0.920 |
| ELW | 0.474 | pass | 0.336 [0.182, 0.528] | 0.147 [0.023, 0.333] | 68 % | 0.916 |
| ERN | 0.508 | pass | 0.346 [0.195, 0.494] | 0.118 [0.016, 0.276] | 70 % | 0.944 |
| EW | 0.340 | pass | 0.472 [0.369, 0.539] | 0.140 [0.008, 0.326] | 83 % | 0.893 |
| KNPE | 0.357 | pass | 0.488 [0.410, 0.546] | 0.213 [0.031, 0.419] | 88 % | 0.843 |
| KNPR | 0.352 | pass | 0.435 [0.294, 0.564] | 0.228 [0.055, 0.451] | 78 % | 0.856 |
| KWPE | 0.307 | pass | 0.492 [0.385, 0.563] | 0.191 [0.008, 0.415] | 84 % | 0.897 |
| LBT | 0.318 | pass | 0.274 [0.163, 0.451] | 0.103 [0.015, 0.248] | 65 % | 0.908 |
| LNAE | 0.465 | pass | 0.292 [0.154, 0.488] | 0.125 [0.023, 0.266] | 62 % | 0.959 |
| LNBE | 0.439 | pass | 0.280 [0.150, 0.467] | 0.096 [0.016, 0.228] | 63 % | 0.958 |
| PH | 0.517 | pass | 0.452 [0.351, 0.516] | 0.125 [0.000, 0.315] | 81 % | 0.904 |
| SAI | 0.371 | pass | 0.372 [0.219, 0.516] | 0.147 [0.023, 0.323] | 75 % | 0.932 |
| STI | 0.469 | pass | 0.371 [0.221, 0.495] | 0.154 [0.016, 0.322] | 76 % | 0.931 |
| TBN | 0.558 | pass | 0.374 [0.252, 0.474] | 0.103 [0.023, 0.214] | 76 % | 0.773 |
| TL | 0.598 | pass | 0.414 [0.295, 0.503] | 0.140 [0.031, 0.295] | 79 % | 0.976 |

Methods, 52 augmented and 17 base (Table S20); environments, five sites; genotypes per environment, the median over environments. Reversal is between the two official metrics, with 95 % intervals. Best base r is the highest mean within-environment Pearson correlation reached by a base method, the quantity of the quality gate;

it never falls below 0.307, so no trait was excluded on that ground. Interval width is the median 95 % rank-interval width as a fraction of the 52-method board. $\Delta r^*$ is not estimable here: the estimator needs more than five environments, and CUBIC has five.

### Section S12. Composition of the constructed method panels

The constructed panels share a core and differ at the margins. Seven base methods of each leave-genotypes-out panel, and eight of the soybean panel, were augmented with the shrunk, inflated and biased variants of Section 2.3; in common bean, spring wheat and Chinese maize the same seven also received one noise-injected and one partially rank-shuffled variant, and in soybean six received noise at two levels (0.5 and 1.5 SD of the predictions) and four a partial rank shuffle. Noise (1 SD of the predictions in the other panels) and the shuffle of 35 % of values are applied to a method's pooled predictions, so they perturb between- as well as within-environment order. This gives the method counts of Table 1. Ridge regressions use an environment index and the first *d* marker principal components as predictors, with an unpenalised intercept, and in soybean also each genotype's training mean. In the leave-genotypes-out panels the environment index of a fold's test genotypes is the mean of that fold's training genotypes in the environment, and the mixed models use that fold's own environment effects, so no test phenotype enters a prediction. The two mixed models are reduced-rank GBLUP models on the same principal components as the ridge regressions; the retained components explain 81.7 % of the marker variance in common bean (80 components), 65.5 % in spring wheat (80), 54.0 % in soybean (100) and 48.6 % in Chinese maize (60); they share one implementation, analysis/crosscrop/code/mixed_models.py, which estimates the variance components by REML after projecting out the environment fixed effects. The learners were run at the conventional settings of Table S20 rather than tuned: the analyses compare metrics applied to one common set of predictions, and the tier partition of the metrics does not depend on how good those predictions are. Scripts: analysis/crosscrop/code/panel_{bean,ursn,nust,china}.py, widen_panel.py (soybean variants).

***Table S20. Base methods and settings in each panel.***

| | **common bean** | **spring wheat** | **soybean** | **Chinese maize** |
|---|---|---|---|---|
| Base methods | 17 | 17 | 26 | 17 |
| Ridge, principal components *d* | 5, 20, 40, 80 | 5, 20, 40, 80 | 10, 25, 50, 100 | 5, 15, 30, 60 |
| Ridge penalties λ | 1, 100 | 1, 100 | 10, 1,000 | 1, 100 |
| Random forest | 300 trees, leaf ≥ 3 | 300 trees, leaf ≥ 3 | 120 trees, depth 8, leaf ≥ 8; 300 trees, leaf ≥ 2 | 300 trees, leaf ≥ 3 |
| Gradient boosting (histogram) | depth 4, 300 iterations, rate 0.06 | same | depth 3, 250, 0.06; depth 8, 500, 0.05 | depth 4, 300 iterations, rate 0.06 |
| *k*-nearest neighbours | *k* = 10, 30 | *k* = 10, 30 | *k* = 10, 50 | *k* = 10 |
| Multilayer perceptron | 48–24 units, early stopping | same | 64–32 units, early stopping | 48–24 units, early stopping |
| Marker-only ridge | *d* = 80 | *d* = 80 | *d* = 25, 100 | — |
| Reduced-rank GBLUP (marker principal | yes | yes | yes | yes |

| | | | | |
|---|---|---|---|---|
| components, REML) | | | | |
| Marker × environment GBLUP (REML) | environment | environment | location | environment |
| Reaction norm | — | — | Finlay–Wilkinson | environmental covariates |
| Other | — | — | genotype mean; genotype-by-environment ridge; environment plus genotype | — |
| Ensembles | one | one | all; linear; tree-based | linear; machine-learning (random forest, gradient boosting, perceptron) |

### Section S13. Which submission stands for a team

The 2022 competition ranked teams on the mean within-environment RMSE, so the leaderboard analyses take each team's best submission under that metric, which reproduces the published leaderboard (analysis/g2f_leaderboard/code/build_inputs.py). That rule is itself one of the two metrics compared, and a rule built on the 2022 metric could favour the 2022 ordering, so Table S21 repeats the leaderboard quantities under four alternatives: the best submission under the 2024 official metric, under the admissible metric, the most recent submission, and no per-team selection at all. Of the 124 submissions in the published table, 116 carry all 22 metrics; only those enter. Script: analysis/g2f_leaderboard/code/selection_rule_sensitivity.py.

***Table S21. The leaderboard quantities under five selection rules.***

| **Rule for which submission stands for a team** | **Entries** | **Reversal, two official metrics** | **Top three by mean_pearson_r** | **1st–2nd gap** | **1st–3rd gap** | **Entries within 0.03 of the leader** |
|---|---|---|---|---|---|---|
| best mean_RMSE (2022 official; used in the paper) | 30 | 26.9 % | CLAC, CGM, UCD_MegaLMM | 0.0033 | 0.0183 | 3 |
| best mean_pearson_r (2024 official) | 30 | 28.7 % | CLAC, CGM, MPB_Group | 0.0033 | 0.0141 | 4 |
| best mean_spearman_r (admissible) | 30 | 32.2 % | CLAC, CGM, MPB_Group | 0.0033 | 0.0141 | 4 |
| most recent submission | 30 | 33.1 % | CLAC, CGM, UCD_MegaLMM | 0.0033 | 0.0193 | 3 |
| every scored | 116 | 28.7 % | CLAC, CGM, | 0.0033 | 0.0141 | 6 |

| | | | |
|---|---|---|---|
| submission, no selection | | | MPB_Group |

The rule used in the paper gives the *lowest* reversal rate of the five (26.9 % against 28.7–33.1 %), so it is the conservative choice for Section 3.2. The first-to-second gap is 0.0033 under every rule. What does move is the third place and the size of the tie at the top: UCD_MegaLMM under the paper's rule, MPB_Group under a rule built on either correlation metric, which lifts the first-to-third gap's counterpart to 0.0141 and puts four entries rather than three within the Gaussian design value of the leader. The statement in Section 3.3 that survives every rule is the first: first and second place are 0.003 apart, an order of magnitude below that value.

### Section S14. Tied predictions and the shortfall below $i_k \cdot r$

Where genotypes share the *k*-th largest predicted value the selected set is not unique (Section S10), and the sort breaks the tie arbitrarily. Because the tied genotypes are then taken in an order unrelated to their observed values, a boundary tie can only lower the realised selection differential, which raises the question whether the shortfall of Section S19 is an artefact of ties. Table S22 answers it by recomputing each verified submission's realised differential under the best and the worst possible tie-break. Script: analysis/crosscrop/code/tie_effect_on_gap.py.

***Table S22. The shortfall below $i_k \cdot r$ under three tie-breaks.***

Mean over the five verified submissions, in phenotypic standard deviations; a positive value means the projection exceeds the realised differential. Environments are counted per submission: 23 each, except 21 for Niche Squad, whose predictions are constant in two environments, so that no selection is defined there; Table S17 counts all 115 cases.

| **Selection fraction *f*** | **Environments with a tie at the boundary (of 113)** | **Shortfall as analysed** | **Shortfall, ties resolved in the selected set's favour** | **Shortfall, ties resolved against it** |
|---|---|---|---|---|
| 0.05 | 0 | 0.3081 | 0.3081 | 0.3081 |
| 0.10 | 2 | 0.1850 | 0.1850 | 0.1860 |
| 0.20 | 4 | 0.1287 | 0.1283 | 0.1290 |

Resolving every tie in the selected set's favour moves the mean shortfall by at most 0.002 SD, against a shortfall of 0.13 to 0.31 SD. Ties therefore do not account for it.

### Section S15. Selection on genotype means across environments

Many multi-environment programmes advance genotypes on their mean over a trial network rather than within each trial. This decision reads the predictions through the ranking of genotype means, so by the construction of Section 2.1 its admissible metrics are those that depend on the predictions only through that ranking — the rank correlation of predicted with observed genotype means is the natural one. None of the 22 official metrics has this form: the mean_ metrics score each environment separately, and the pooled metrics score cells rather than genotype means.

The outcome test of Section 2.4 was repeated for this decision (Table S23). Within each environment, predictions and observations were centred (removing environment main effects) and observations divided by their standard deviation; a genotype's mean was taken over the environments of a half in which it was tested at least twice. In half B the top 10 % of eligible genotypes by predicted mean were advanced, and the outcome is their mean observed value minus that of all eligible genotypes. Each rule ranked the methods on half A — the within-environment rules as in Table S7, the two genotype-mean rules on half-A genotype means — and recovery is normalised as in Table S7. Point estimates average 300 random splits; intervals resample environments (200 replicates × 25 splits of the distinct resampled environments). Splits in which a genotype-mean rule could not be scored (no genotype tested twice in a small resampled half) are skipped for that rule. Script: analysis/crosscrop/code/decision_genotype_means.py.

***Table S23. Outcome test for selection on genotype means across environments.***

Selection gain recovered (%) [95 % CI]; paired rows are differences in percentage points. Datasets whose outcome reliability falls below 0.5 are reported but not compared (Section 2.4 of the main text); five maize submissions are too few for a reliability estimate (—).

| **Rule** | **maize (5 submissions)** | **common bean** | **spring wheat** | **soybean** |
|---|---|---|---|---|
| Genotype-mean rank correlation (admissible for this decision) | 60.3 [−44.6, 83.2] | 62.8 [41.2, 76.0] | 34.9 [−0.4, 58.1] | 85.8 [63.2, 95.0] |
| Genotype-mean Pearson correlation | 11.6 [−25.7, 48.5] | 61.8 [35.5, 75.9] | 52.7 [6.1, 66.3] | 84.9 [62.2, 95.7] |
| mean_spearman_r | 60.8 [−35.3, 85.3] | 51.2 [37.7, 75.2] | 49.6 [15.5, 63.3] | 84.7 [58.8, 95.0] |
| mean_pearson_r | 11.6 [−25.7, 47.3] | 49.6 [36.8, 74.5] | 41.6 [1.9, 76.2] | 84.5 [57.3, 96.1] |
| mean_NDCG@10% | 58.2 [7.8, 86.7] | 67.9 [32.9, 84.2] | 13.6 [−47.6, 47.0] | 84.2 [44.9, 95.9] |
| mean_RMSE | −28.1 [−109.5, 54.3] | 53.7 [36.5, 74.6] | 62.6 [15.9, 70.7] | 44.3 [8.8, 79.3] |
| mean_MAE | −28.4 [−100.0, 54.3] | 50.9 [36.7, 74.8] | 64.3 [15.6, 75.7] | 33.3 [4.5, 71.2] |
| mean_r2_score | −95.9 [−130.2, 38.8] | 59.2 [36.4, 75.5] | 40.0 [−5.2, 71.6] | 23.1 [−3.9, 54.0] |
| Paired: mean_spearman_r − genotype-mean rank correlation | 0.5 [−29.3, 48.4] | −11.5 [−16.8, 8.5] | 14.7 [−14.3, 41.1] | −1.2 [−13.1, 6.5] |
| Paired: mean_spearman_r − mean_RMSE | 88.9 [−36.7, 168.8] | −2.4 [−11.9, 11.0] | −13.0 [−35.6, 26.9] | 40.4 [1.4, 73.7] |
| Paired: genotype-mean rank correlation − mean_RMSE | 88.4 [−48.1, 172.0] | 9.1 [−9.3, 22.1] | −27.7 [−48.7, 10.5] | 41.6 [2.7, 73.1] |
| Outcome reliability | — | 0.780 | 0.467 | 0.668 |
| Methods / environments / genotypes | 5 / 23 / 548 | 52 / 10 / 467 | 52 / 109 / 384 | 66 / 169 / 1,161 |

Where the outcome is reliable, in common bean and soybean, the within-environment rank correlation recommended in the main text shows no detectable difference from the decision's own admissible metric: the two differ by −11.5 [−16.8, 8.5] percentage points in common bean and −1.2 [−13.1, 6.5] in soybean. In soybean both lead mean_RMSE by about 40 points, with intervals excluding zero; in common bean mean_RMSE does about as well as either, with both paired intervals including zero. These intervals do not establish equivalence; with the present number of environments, differences of about 15 percentage points cannot be excluded. In spring wheat the outcome of this decision orders methods with a split-half reliability of 0.467, below the 0.5 criterion, so its column is reported but not compared. Admissibility for this decision does not make the genotype-mean correlation a better selector of methods than the within-environment one — its genotype means are estimated from half the environments and are noisier — but it does make it the metric that cannot be moved without moving the decision.

### Section S16. Withdrawn panels and exploratory analyses

Two further panels were built on the same design and withdrawn: rice, from 107 IRRI elite breeding lines grown in 15 environments in Asia and Africa (grain yield) [S8], and wheat, from 121 CAIGE lines in 13 site-years (grain yield) [S9]. In both, every method ranks genotypes within an environment at or near chance: the best mean within-environment Pearson correlation is 0.081 among the 51 rice methods, 24 of which are negative, and 0.114 among the 44 wheat methods, 10 of which are negative. A panel in which no method ranks genotypes better than chance cannot test claims about ranking methods — its reversal rates, rank ranges and thresholds are driven by noise in the metrics being compared — so both are excluded from every cross-dataset analysis by analysis/crosscrop/code/scope.py. They were built with an earlier version of the panel code, before GBLUP was added and before the correction of the environment index described in Section S12, and have not been rebuilt. Their scripts (panel_rice.py, panel_wheat.py) and the rice primary files are kept in the repository for the record; the CAIGE compilations carry no redistribution licence and are not included.

*Two exploratory analyses, not carried into the paper,* are kept in the repository (analysis/easygese/). First, EasyGeSe [S10] benchmarks ten genomic prediction models on 93 traits in 10 species by pooled cross-validation; there, Pearson $r$ and RMSE reverse 9.1 % of model pairs (95 % CI 8.2–10.0 %), and 80 of the 93 traits show at least one reversal. EasyGeSe records no environments, so the within-environment decision cannot be evaluated on it. Second, averaging each genotype over environments before scoring — a pooled design — changes the reversal rate between the same two metrics from 46.7 % to 37.6 % in spring wheat, from 39.3 % to 40.6 % in common bean and from 34.3 % to 42.1 % in soybean, so disagreement between these metrics in the panels does not depend on scoring within environments; the collapsed design scores a different decision, selection on genotype means, treated in Section S15.

### Section S17. Panel composition and the outcome test

The two mixed models were added to the panels after the other methods, and in soybean they are the methods the correlation rules select. Table S24 therefore repeats the outcome test of Table S7 without them, and with each base method of the base panels left out in turn (together with its variants). Recovery is computed as in Table S7; the full-panel rows reproduce Table S7 exactly. Script: analysis/crosscrop/code/composition_sensitivity.py.

***Table S24. Outcome test under changes to the composition of the method panels.***

Selection gain recovered by the first choice of each family's rules (range over the family: correlation, mean_pearson_r and mean_spearman_r; error magnitude, mean_RMSE, mean_MAE and mean_r2_score), 400 environment splits.

| **Dataset** | **Panel** | **Methods removed** | **Methods** | **Correlation family** | **Error-magnitude family** | **mean_NDCG@10%** |
|---|---|---|---|---|---|---|
| soybean | augmented | none | 66 | 98 to 99 % | 13 to 34 % | 99 % |
| soybean | augmented | the two GBLUP models | 64 | 78 to 80 % | 27 to 31 % | 47 % |
| soybean | base | none | 26 | 97 to 99 % | −6 to 19 % | 99 % |
| soybean | base | the two GBLUP models | 24 | 68 to 71 % | −6 to 0 % | 23 % |

Leave-one-method-out, base panels, 200 splits per panel: the number of panels in which both correlation rules recover more than mean_RMSE, and the range of the difference in percentage points.

| **Dataset** | **Panels** | **Both correlation rules above mean_RMSE** | **mean_pearson_r − mean_RMSE** | **mean_spearman_r − mean_RMSE** |
|---|---|---|---|---|
| soybean | 26 | 26 of 26 | 49 to 82 | 46 to 78 |

Without the GBLUP models the correlation family still leads in soybean, and with one base method left out the soybean result holds in every panel.

### Section S18. GPverdict: implementation, validation and the CUBIC verdict

GPverdict 1.1.1 (https://doi.org/10.5281/zenodo.23051854; source https://github.com/nblvguohao/gpverdict; web version https://nblvguohao.github.io/gpverdict/) implements Sections 2.1–2.4 of the main text for any table of cross-validated predictions with columns environment, genotype, observed value, prediction and method. It needs only numpy and pandas, so the same code runs as a Python package, from the command line and, through Pyodide, inside a web browser, where the data are processed on the user's computer and never uploaded. For each method and environment it computes eleven metrics — the rank correlation, realised selection differential, top-*k* hit rate and NDCG (Tier I by construction), Pearson *r* (Tier II), and RMSE, MAE, $R^2$_score, the calibration slope and the pooled Pearson *r* and RMSE (Tier III) — and checks their tiers empirically on the user's predictions with the affine, monotone and reversal tests of Section 2.4 (three replicates each on up to five methods). Methods are ranked by the mean within-environment rank and Pearson correlations, and a method counts as tied with the leaders if it lies within the resolvable gap of the best on either, with 95 % rank intervals from 400 environment resamples when there are at least eight environments. The resolvable gap is $k(f)/\sqrt{N}$, with *N* the median number of genotype–environment cells per method and *k* interpolated log-linearly between the design-grid constants 4.29, 2.97 and 2.09 at $f$ = 0.05, 0.10 and 0.20 (Table S11); the report also gives the 1.5–4.2-fold range that the calibrated panels needed (Table S10). The outcome test is that of Section 2.4 with 200 splits; with fewer than eight environments, genotypes are split in halves within each environment instead. The report states the verdict in plain language and gives the cells needed to resolve accuracy gains of 0.01–0.10. Fig. S5 shows the web version running its bundled spring wheat example.

Table S26 shows that GPverdict reproduces the published values on the spring wheat base panel exactly; the package's tests check the same values (tests/test_reproduces_paper.py), and these spring wheat predictions are its example data set. Table S25 is the CUBIC report summarised in Section 3.6 of the main text. Script: analysis/crosscrop/code/gpverdict_applications.py.

***Table S25. GPverdict on CUBIC ear weight: the 17 base methods ranked by the admissible metric.***

Mean over the five sites of each within-environment statistic; realised selection differential of the top 10 % in phenotypic SD. † Within 0.036 of the leader, the smallest gap this trial resolves (6,699 cells per method; Gaussian lower bound, calibrated panels needed 0.054–0.152). Method codes as in Table S20: ridge_pc*d*_lo/_hi, ridge regression on *d* principal components with penalty 1 or 100; rf, random forest; gbm, gradient boosting; mlp, multilayer perceptron; knn10, *k*-nearest neighbours; reaction_norm_EC, reaction norm on environmental covariates; gblup, gblup_gxe, the two mixed models; ens_lin, ens_ml, the linear and the machine-learning ensemble.

| **Rank** | **Method** | **Rank correlation** | **Pearson *r*** | **RMSE** | **Realised selection differential** | **Rank by RMSE** |
|---|---|---|---|---|---|---|
| 1 | ens_ml † | 0.341 | 0.339 | 26.28 | 0.573 | 2 |
| 2 | rf † | 0.340 | 0.340 | 26.21 | 0.624 | 1 |
| 3 | gbm † | 0.323 | 0.324 | 26.47 | 0.592 | 3 |
| 4 | gblup_gxe | 0.287 | 0.290 | 26.66 | 0.501 | 4 |
| 5 | ridge_pc60_lo | 0.267 | 0.269 | 26.92 | 0.482 | 8 |
| 6 | ridge_pc60_hi | 0.267 | 0.269 | 26.92 | 0.482 | 7 |
| 7 | gblup | 0.267 | 0.270 | 26.85 | 0.492 | 5 |
| 8 | knn10 | 0.259 | 0.263 | 31.40 | 0.547 | 17 |
| 9 | mlp | 0.258 | 0.253 | 28.88 | 0.413 | 16 |
| 10 | ens_lin | 0.255 | 0.260 | 26.90 | 0.534 | 6 |
| 11 | reaction_norm_EC | 0.233 | 0.237 | 27.08 | 0.536 | 9 |
| 12 | ridge_pc30_lo | 0.229 | 0.233 | 27.13 | 0.512 | 11 |
| 13 | ridge_pc30_hi | 0.229 | 0.233 | 27.13 | 0.512 | 10 |
| 14 | ridge_pc15_lo | 0.216 | 0.217 | 27.20 | 0.484 | 13 |
| 15 | ridge_pc15_hi | 0.216 | 0.217 | 27.20 | 0.484 | 12 |
| 16 | ridge_pc5_hi | 0.043 | 0.072 | 27.77 | 0.145 | 14 |
| 17 | ridge_pc5_lo | 0.043 | 0.072 | 27.77 | 0.145 | 15 |

Out of sample, splitting lines in halves within each site (200 splits; outcome reliability 0.80), the method each rule picks recovers 78.8 % (rank correlation), 82.8 % (Pearson *r*) and 90.0 % (RMSE) of the selection gain attainable over an average method. RMSE and Pearson *r* order 14.0 % of the 136 method pairs in opposite directions.

***Table S26. GPverdict against the published values (spring wheat, base panel, 17 methods).***

| Quantity | GPverdict | Published (Tables S4, S7) | Absolute difference |
|---|---|---|---|
| reversal rate, mean_RMSE vs mean_pearson_r | 0.2500 | 0.2500 | 0e+00 |
| outcome-test recovery, spearman | 0.5872 | 0.5872 | 0e+00 |
| outcome-test recovery, pearson | 0.7845 | 0.7845 | 0e+00 |
| outcome-test recovery, rmse | 0.3790 | 0.3790 | 0e+00 |

The values above are those of the Python package at its default 400 splits; the browser version caps the bootstrap at 200 splits and reports whole percentages, so its recoveries differ from these by at most 1.5 percentage points.

### Section S19. Methods and results details for the main text

Details summarised in the main text, given here in full; the section each paragraph supports is named in its heading.

**Maize submissions (Section 2.2).** As on the official leaderboard, a team is represented by its best submission under the 2022 official metric; because that rule is itself one of the two metrics compared, Table S21 repeats the leaderboard results under four alternatives. Seven submission files were recoverable from the team repositories named in the paper's data-availability statement; five reproduce the official values to within $5 \times 10^{-7}$, the precision of the six decimals published, and are used wherever per-genotype predictions are required. They come from three teams — three from Kernel of Truth (244906, 244944, 244953), one each from EnBiSys (243568, that team's leaderboard entry) and Niche Squad (244985). Two files match none of their team's scored submissions and were excluded as post-hoc versions.

**Metric panel and prediction panels (Section 2.3).** Every panel contains ridge regression on an environment index and marker principal components at four dimensionalities crossed with two penalties, random forests, histogram gradient boosting, *k*-nearest neighbours, a multilayer perceptron and unweighted ensembles, together with the two mixed models most used in multi-environment genomic prediction, built on the same marker principal components: a reduced-rank GBLUP (environment fixed effects plus a genomic random effect on those components) and its marker × environment extension (each environment — each location in the forward-year soybean panel — carrying its own genomic effect), both fitted by restricted maximum likelihood. All three add a marker-only ridge using no phenotype-derived feature, and the soybean panel a genomic Finlay–Wilkinson reaction norm and genotype-mean and genotype-by-environment ridge models. Marker matrices were centred, scaled and reduced by singular value decomposition, retaining 80–100 principal components depending on dataset; genotype identifiers were matched to marker files after stripping non-alphanumeric characters, which in soybean recovers all 2,666 raw names (2,513 genotypes after normalisation) against 615 by raw matching. In the leave-genotypes-out panels a fold's test genotypes take the environment index and mixed-model environment effects estimated from that fold's training genotypes, so no test phenotype enters a prediction.

Panels fitted by least squares are homogeneous in calibration in a way that real competition entries are not: the G2F leaderboard spans mean RMSE 2.33–10.47 (CV = 0.49) while an unmodified soybean panel spans 12.5–14.1 (CV = 0.02).

**Statistical procedures (Section 2.4).** *Reversal tests.* For the 231 metric pairs in maize, each rate was tested against $H_0$: rate ≤ 0.05 by a one-sided Wald test with a team-bootstrap standard error (4,000 replicates), with Benjamini–Hochberg control across pairs.

*Realised selection differential.* $k$ is $fn$ rounded to the nearest integer; ties in the predictions are broken by the sort, arbitrarily but deterministically. $i_k \cdot r$ is the selection-differential identity behind the breeder's equation, here in phenotypic units. The copula benchmark carries each environment's observed yield distribution and rank correlation.

*Threshold.* The threshold is reported as not reached when it exceeds the largest observed $\Delta r$. An isotonic fit checks the logistic shape, and an out-of-sample variant estimates $\Delta r$ and $\Delta g$ on disjoint halves of the environments. The Gaussian panels are simulated at stated numbers of methods, genotypes per environment and environments, accuracy range and selection fraction; method $m$ predicts $r_m y + \sqrt{(1 - r_m^2)}\, e_m$, and the errors share a fraction $\lambda$ of their variance.

*Heritability ceiling.* Components were estimated per environment by unbalanced one-way ANOVA [S2,S3] on the 192 environments of the G2F 2014–2021 training data that pass the retention rules of Section S6 (114,234 of its 128,167 plots), and for common bean from raw plots and from published BLUE standard errors as $1 - \text{mean}(SE^2)/\text{var}(BLUE)$.

**Rank intervals (Section 3.2).** Removing redundant metrics leaves the maize width at 17 places (Table S8).

**Shortfall below $i_k \cdot r$ (Section 2.4).** For the five verified G2F submissions, the realised selection differential of each method's own top-$k$ pick falls below $i_k \cdot r$ in all 15 comparisons: by a mean of 0.31, 0.19 and 0.13 phenotypic standard deviations at $f = 0.05$, 0.10 and 0.20, and at $f = 0.10$ by 0.095, 0.240, 0.110, 0.309 and 0.172 SD. Standard errors over the 21–23 environments are 0.019–0.046 SD, $|t|$ is 2.5–10.5, and $P \leq 0.001$ in 12 of the 15; one submission's top decile performed below its environment mean. These five submissions come from three teams that the 2022 leaderboard placed 21st, 25th and 28th of 30, and are the only files that could be recovered and verified against the published metrics (Section 2.2 of the main text), so the size of the shortfall is theirs rather than an estimate for the competition. The choice of correlation coefficient accounts for part of the shortfall, not most, and the shape of the yield distribution for almost none. Against a Gaussian copula carrying each environment's observed yield distribution and its observed rank correlation, the mean shortfall over $f = 0.10$ and 0.20 falls from 0.156 against the finite-population benchmark to 0.098 SD but stays negative for every submission at every intensity, with residual $|t|$ of 2.5–8.2; the copula mainly corrects Pearson $r$'s overstatement of rank agreement, and the remainder is non-Gaussian dependence (Table S9). Nor is the shortfall a finite-population artefact — with a median of 445 genotypes per environment, replacing $i_k$ by the finite-population expectation changes the projection in the third decimal and explains none of the gap — or an effect of tied predictions: resolving every tie in the selected set's favour moves the mean shortfall by at most 0.002 SD (Table S22). The predictions therefore carry less information about the top of the distribution than their overall correlation implies. A related result is known for breeding values: accuracies estimated in an unselected reference population do not carry over to the selected fraction [S11]. Breeding programmes are now asked to report realised genetic gain across breeding cycles [S12]; setting realised selection against its projection, as here, is the within-cycle counterpart.

**Threshold (Section 3.3).** Among base-method pairs with $\Delta r < 0.01$ the probability is 50 % and 67 % in common bean and soybean (spring wheat has seven such pairs); even below $\Delta r = 0.01$, 24–36 % of the two methods' shortlists differ in those two panels.

Estimating $\Delta r$ and $\Delta g$ on disjoint halves of the environments gives 0.126, 0.253 and 0.179, and relaxing selection to $f = 0.20$ gives 0.037, 0.116 and 0.042 (Table S5).

**Rank-interval transfer (Section 3.2).** The interval transfers when one metric is left out (coverage 94.2 %) but not across classes: calibrated on the error-magnitude metrics, it covers the correlation metrics' ranks 56 % of the time.

### Section S20. Pooled metrics decomposed into an environment-mean part and a within-environment part

**Set-up.** Environment $j$ holds $n_j$ genotype–environment cells, $N = \Sigma\, n_j$ and $w_j = n_j / N$; means, variances and covariances are taken over cells. Write every prediction as $\hat{y}_{ij} = m_j + d_{ij}$, with $m_j$ the mean prediction in environment $j$ and $\Sigma_i\, d_{ij} = 0$ in every environment. This is notation, not a model: any set of predictions can be written so. Define $a = \Sigma\, w_j\, (m_j - \bar{m})(\bar{y}_j - \bar{y})$, $v\_b = \Sigma\, w_j\, (m_j - \bar{m})^2$, $u^2 = \Sigma\, w_j\, \mathrm{Var}_j(d)$, $k = \Sigma\, w_j\, \mathrm{Cov}_j(d, y) / u$ and $S = \mathrm{Var}(y)$.

**Proposition 1.** Pooled Pearson correlation is $r(u) = (a + k\,u) / \sqrt{((v\_b + u^2)\, S)}$. *Proof.* Because $d$ has mean zero within every environment and $m$ is constant within it, the between–within decomposition of the covariance and the variance has no cross terms: $\mathrm{Cov}(\hat{y}, y) = a + k\,u$ and $\mathrm{Var}(\hat{y}) = v\_b + u^2$. No equality of environment sizes or variances, and no independence of $m$ and $d$, is needed. ■

**Proposition 2.** If $a > 0$ and $k > 0$, $r(u)$ has a unique maximum over $u \geq 0$ at $u^* = k\,v\_b / a$, where $r^2\_max = (a^2/v\_b + k^2)/S$. *Proof.* $r'(u)$ is proportional to $k\,v\_b - a\,u$, positive below $u^*$ and negative above it; substituting gives $r\_max$. ■ The share of $r^2\_max$ carried by the environment means is $\varphi = (a^2/v\_b) / (a^2/v\_b + k^2)$. Where $\varphi$ is near one, pooled correlation is nearly insensitive to $k$, the within-environment information that selection uses, and rewards instead a scale $u$ close to $u^*$.

**Proposition 3.** Pooled mean squared error is $\Sigma\, w_j\, (m_j - \bar{y}_j)^2 + u^2 - 2\,k\,u + \Sigma\, w_j\, \mathrm{Var}_j(y)$, minimised at $u = k$ whatever $a$. Its first term depends on the environment means alone. ■

Within-environment Pearson and rank correlations depend on neither $m_j$ nor $u$, which is the content of admissibility (Tier II and Tier I).

**Numerical check** (500 random designs with 3–39 environments of 3–299 cells, unequal within-environment variances and profile scales, environment means correlated with the profile; analysis/supplementary/code/pooled_decomposition.py): cases with a > 0 and k > 0: 388 of 500; max |r_numeric - r_formula|: 2.00e-15; max relative error of u* beyond one grid step: 0.00e+00; max |max r_numeric - r_max|: 2.46e-08; max |argmin MSE - k| beyond one grid step: 0.00e+00.

**The paper's predictions.** The identity reproduces pooled *r* for every method to machine precision. Table S27 summarises the components by dataset: the five verified G2F submissions and every base method of the three panels.

***Table S27. Pooled Pearson correlation and pooled MSE decomposed.***

| Dataset | Methods | Pooled r, median [range] | Within-environment r, median [range] | φ, median [range] | Share of pooled MSE in the environment-mean term, median [range] |
|---|---|---|---|---|---|
| common bean | 17 | 0.85 [0.22, 0.86] | 0.40 [0.37, 0.45] | 0.94 [0.35, 0.95] | 0.01 [0.00, 0.69] |
| maize (5 verified submissions) | 5 | 0.13 [-0.07, 0.50] | 0.13 [0.10, 0.22] | 0.70 [0.08, 0.99] | 0.64 [0.59, 0.69] |
| soybean | 26 | 0.45 [0.18, 0.50] | 0.17 [0.15, 0.36] | 0.95 [0.65, 0.98] | 0.63 [0.56, 0.70] |
| spring wheat | 17 | 0.71 [0.15, 0.75] | 0.16 [-0.08, 0.40] | 0.95 [0.66, 1.00] | 0.02 [0.01, 0.43] |

In the three panels φ is 0.94–0.95 at the median, and pooled *r* is two to four times the within-environment *r* in common bean and spring wheat. The share of pooled MSE in the environment-mean term follows the design: 0.01–0.02 at the median in the two leave-genotypes-out panels, whose test environments are seen in training, and 0.63 in the forward-in-time soybean panel, whose test years are not; 0.64 across the five competition submissions.

### Section S21. Genetic architecture and the disagreement between metrics

**Data and panels.** Seven public multi-environment datasets were assembled with one design, the Atlas (analysis/atlas/): 23 CUBIC maize traits, six DROPS maize traits, four common bean (VEF) traits, two spring wheat (URSN) traits, ten soybean NUST traits and seven SoyNAM traits, each with a panel of 17 base methods built as in Section 2.3 but without the miscalibration variants (five-fold leave-genotypes-out cross-validation; ridge on an environment index and marker principal components at four dimensionalities and two penalties; marker-only ridge; random forest; gradient boosting; two *k*-nearest-neighbour learners; a multilayer perceptron; GBLUP; marker × environment GBLUP; an unweighted ensemble). Environments with at least 25 genotypes enter (12 in URSN); lodging and disease scores are negated so that the higher value is selected. The G2F entry of the Atlas carries five competition submissions rather than a panel and has no markers, so 52 combinations enter here. CUBIC phenotypes are CC BY-NC-ND, so only summary results are published.

**Plan.** The analysis was specified before the parameters were computed (analysis/genetics/analysis_plan.md): three predictors, one primary outcome, the within-dataset test and Holm correction, the minimum detectable association ($|\rho_w| \approx 0.40$ at 80 % power), and the rule that a null result is reported in text and table without a figure.

**Genetic parameters.** For each combination the marker × environment GBLUP of the panels was fitted once, by REML, to all the data, with environments as fixed effects and the Atlas's marker principal components as features (60 for CUBIC, 80 otherwise). With $d_0 = \sigma^2_u/\sigma^2_e$ and $d_1 = \sigma^2_v/\sigma^2_e$ the genomic main-effect and

G×E variance ratios, the within-environment genomic heritability is $(d_0 + d_1)/(d_0 + d_1 + 1)$, the G×E share is $d_1/(d_0 + d_1)$, and the between-environment genetic correlation of this compound-symmetry model is $r\_g = 1 −$ G×E share. The median between-environment correlation of genotype values (environment pairs sharing at least 25 genotypes) is a model-free companion that mixes heritability and $r\_g$. A non-genetic covariate, the standard deviation across the 17 methods of their mean within-environment calibration slope, asks whether any association merely reflects calibration differences. The parameters of every combination are listed in Table S28.

**Result.** No primary association survives: within datasets, the reversal rate correlates −0.24 with genomic heritability (Holm $P = 0.42$), +0.29 with G×E share (Holm $P = 0.40$), −0.17 with between-environment r (Holm $P = 0.42$) (Table S29). The secondary outcomes show no association either. The one secondary result with a small uncorrected $P$ is the calibration spread itself, in the direction opposite to intuition (more spread, fewer reversals); it is reported, not interpreted. The environment counts of three NUST and SoyNAM combinations exceed those in the Atlas table, because GPverdict's scoring drops a few environments that the loader keeps; the reversal rates are the Atlas's.

***Table S28. Genetic parameters of the 52 combinations.***

| Crop | Dataset | Trait | Env. | Genotypes | h²_g | G×E share | r_g | Between-env. r | Calibration SD | RMSE–Pearson reversal (%) |
|---|---|---|---|---|---|---|---|---|---|---|
| maize | CUBIC | ATI | 5 | 1,402 | 0.15 | 0.14 | 0.86 | 0.33 | 0.22 | 23 |
| maize | CUBIC | CW | 5 | 1,401 | 0.20 | 0.20 | 0.80 | 0.52 | 0.18 | 19 |
| maize | CUBIC | DTA | 5 | 1,402 | 0.24 | 0.14 | 0.86 | 0.53 | 0.24 | 18 |
| maize | CUBIC | DTS | 5 | 1,402 | 0.20 | 0.15 | 0.85 | 0.54 | 0.22 | 20 |
| maize | CUBIC | DTT | 5 | 1,402 | 0.24 | 0.15 | 0.85 | 0.56 | 0.22 | 20 |
| maize | CUBIC | ED | 5 | 1,402 | 0.15 | 0.06 | 0.94 | 0.40 | 0.23 | 18 |
| maize | CUBIC | EH | 5 | 1,403 | 0.25 | 0.02 | 0.98 | 0.67 | 0.17 | 26 |
| maize | CUBIC | EL | 5 | 1,403 | 0.19 | 0.07 | 0.93 | 0.56 | 0.23 | 21 |
| maize | CUBIC | ELL | 5 | 1,403 | 0.21 | 0.10 | 0.90 | 0.63 | 0.16 | 23 |
| maize | CUBIC | ELW | 5 | 1,403 | 0.22 | 0.05 | 0.95 | 0.58 | 0.23 | 24 |
| maize | CUBIC | ERN | 5 | 1,403 | 0.21 | 0.05 | 0.95 | 0.58 | 0.21 | 20 |
| maize | CUBIC | EW | 5 | 1,400 | 0.12 | 0.19 | 0.81 | 0.41 | 0.16 | 24 |
| maize | CUBIC | KNPE | 5 | 1,403 | 0.12 | 0.15 | 0.85 | 0.45 | 0.13 | 28 |
| maize | CUBIC | KNPR | 5 | 1,403 | 0.49 | 0.02 | 0.98 | 0.44 | 0.19 | 27 |
| maize | CUBIC | KWPE | 5 | 1,387 | 0.10 | 0.22 | 0.78 | 0.37 | 0.16 | 29 |
| maize | CUBIC | LBT | 5 | 1,403 | 0.13 | 0.11 | 0.89 | 0.37 | 0.23 | 15 |
| maize | CUBIC | LNAE | 5 | 1,403 | 0.22 | 0.02 | 0.98 | 0.51 | 0.24 | 18 |
| maize | CUBIC | LNBE | 5 | 1,403 | 0.22 | 0.02 | 0.98 | 0.54 | 0.24 | 14 |
| maize | CUBIC | PH | 5 | 1,403 | 0.30 | 0.01 | 0.99 | 0.70 | 0.14 | 23 |
| maize | CUBIC | SAI | 5 | 1,402 | 0.14 | 0.08 | 0.92 | 0.42 | 0.23 | 22 |

| | | | | | | | | | | |
|---|---|---|---|---|---|---|---|---|---|---|
| maize | CUBIC | STI | 5 | 1,402 | 0.20 | 0.06 | 0.94 | 0.48 | 0.22 | 15 |
| maize | CUBIC | TBN | 5 | 1,403 | 0.27 | 0.01 | 0.99 | 0.70 | 0.18 | 17 |
| maize | CUBIC | TL | 5 | 1,403 | 0.32 | 0.01 | 0.99 | 0.72 | 0.17 | 15 |
| common bean | VEF | seed yield | 10 | 467 | 0.31 | 0.40 | 0.60 | 0.34 | 0.15 | 16 |
| common bean | VEF | 100-seed weight | 9 | 443 | 0.38 | 0.12 | 0.88 | 0.53 | 0.19 | 16 |
| common bean | VEF | days to flowering | 8 | 467 | 0.39 | 0.36 | 0.64 | 0.44 | 0.26 | 20 |
| common bean | VEF | days to physiological maturity | 6 | 467 | 0.39 | 0.63 | 0.37 | 0.24 | 0.33 | 14 |
| wheat | URSN | FHB resistance (100 - disease score) | 109 | 384 | 0.57 | 0.54 | 0.46 | 0.59 | 0.30 | 25 |
| wheat | URSN | 100 - visually scabby kernels (%) | 109 | 387 | 0.64 | 0.57 | 0.43 | 0.69 | 0.21 | 35 |
| soybean | NUST | YieldBuA | 548 | 2,512 | 0.43 | 0.75 | 0.25 | 0.25 | 0.20 | 21 |
| soybean | NUST | SeedSize | 499 | 2,512 | 0.48 | 0.37 | 0.63 | 0.72 | 0.23 | 20 |
| soybean | NUST | Maturity | 492 | 2,512 | 0.57 | 0.52 | 0.48 | 0.73 | 0.26 | 26 |
| soybean | NUST | Lodging | 487 | 2,512 | 0.30 | 0.75 | 0.25 | 0.34 | 0.28 | 36 |
| soybean | NUST | Height | 467 | 2,510 | 0.40 | 0.55 | 0.45 | 0.56 | 0.21 | 41 |
| soybean | NUST | Protein | 333 | 2,482 | 0.44 | 0.39 | 0.61 | 0.64 | 0.19 | 23 |
| soybean | NUST | Oil | 333 | 2,482 | 0.39 | 0.49 | 0.51 | 0.62 | 0.27 | 12 |
| soybean | NUST | SeedQuality | 448 | 2,512 | 0.23 | 0.90 | 0.10 | 0.08 | 0.45 | 42 |
| soybean | NUST | Chlorosis | 38 | 1,381 | 0.26 | 0.56 | 0.44 | 0.29 | 0.23 | 38 |
| soybean | NUST | SDS | 16 | 423 | 0.37 | 0.67 | 0.33 | 0.25 | 0.14 | 37 |
| maize | DROPS | yield | 10 | 246 | 0.47 | 0.31 | 0.69 | 0.46 | 0.25 | 20 |
| maize | DROPS | grain_number | 10 | 246 | 0.44 | 0.29 | 0.71 | 0.39 | 0.24 | 34 |

| | | | | | | | | | | |
|---|---|---|---|---|---|---|---|---|---|---|
| maize | DROPS | grain_weight_mg | 9 | 246 | 0.38 | 0.13 | 0.87 | 0.56 | 0.20 | 20 |
| maize | DROPS | anthesis_d20C | 10 | 246 | 0.60 | 0.03 | 0.97 | 0.74 | 0.26 | 15 |
| maize | DROPS | silking_d20C | 10 | 246 | 0.59 | 0.03 | 0.97 | 0.71 | 0.24 | 16 |
| maize | DROPS | plant_height_cm | 10 | 246 | 0.28 | 0.07 | 0.93 | 0.41 | 0.23 | 30 |
| soybean | SoyNAM | yield | 18 | 5,182 | 0.29 | 0.51 | 0.49 | 0.23 | 0.22 | 10 |
| soybean | SoyNAM | days_to_maturity | 15 | 5,182 | 0.31 | 0.28 | 0.72 | 0.64 | 0.30 | 24 |
| soybean | SoyNAM | lodging | 15 | 5,182 | 0.27 | 0.51 | 0.49 | 0.30 | 0.25 | 26 |
| soybean | SoyNAM | plant_height_cm | 15 | 5,182 | 0.33 | 0.17 | 0.83 | 0.59 | 0.31 | 26 |
| soybean | SoyNAM | protein | 10 | 5,182 | 0.38 | 0.21 | 0.79 | 0.51 | 0.28 | 23 |
| soybean | SoyNAM | oil | 10 | 5,182 | 0.36 | 0.23 | 0.77 | 0.53 | 0.31 | 15 |
| soybean | SoyNAM | seed_weight_100_g | 10 | 5,182 | 0.42 | 0.17 | 0.83 | 0.61 | 0.33 | 25 |

***Table S29. Tests of the pre-specified plan (10,000 permutations within datasets).***

| **Outcome** | **Predictor** | **Model** | ***n*** | **ρ_w or β** | **Permutation *P*** | **Holm *P*** |
|---|---|---|---|---|---|---|
| RMSE–Pearson reversal | genomic heritability | within-dataset rank correlation | 52 | -0.24 | 0.209 | 0.42 |
| RMSE–Pearson reversal | G×E share | within-dataset rank correlation | 52 | +0.29 | 0.132 | 0.40 |
| RMSE–Pearson reversal | between-environment r | within-dataset rank correlation | 52 | -0.17 | 0.375 | 0.42 |
| methods tied with the leader | genomic heritability | within-dataset rank correlation | 52 | -0.01 | 0.958 | |
| methods tied with the leader | G×E share | within-dataset rank correlation | 52 | -0.10 | 0.616 | |
| methods tied with the leader | between-environment r | within-dataset rank correlation | 52 | +0.03 | 0.893 | |
| gain recovered, correlation − RMSE pick | genomic heritability | within-dataset rank correlation | 51 | +0.27 | 0.180 | |

| | | | | | |
|---|---|---|---|---|---|
| gain recovered, correlation − RMSE pick | G×E share | within-dataset rank correlation | 51 | +0.02 | 0.939 |
| gain recovered, correlation − RMSE pick | between-environment r | within-dataset rank correlation | 51 | +0.27 | 0.185 |
| RMSE–Pearson reversal | genomic heritability | rank regression with calibration SD | 52 | -0.24 | 0.162 |
| RMSE–Pearson reversal | G×E share | rank regression with calibration SD | 52 | +0.26 | 0.121 |
| RMSE–Pearson reversal | between-environment r | rank regression with calibration SD | 52 | -0.26 | 0.126 |
| RMSE–Pearson reversal | calibration SD | within-dataset rank correlation | 52 | -0.46 | 0.012 |

Within the Atlas, therefore, how often RMSE and Pearson correlation disagree is not predicted by heritability, by the G×E share or by the between-environment correlation. A programme cannot infer from the genetic parameters of its trait that the check is unnecessary; it has to be run on the programme's own predictions.

### Section S22. The resolution threshold and tests that compare two correlations

The pairs are those of the threshold analysis (base methods, $f = 0.10$, at least eight shared environments, ties dropped): 133 in common bean, 132 in spring wheat and 324 in soybean. For each pair three tests of the accuracy difference were run, two-sided at 0.05. *Meng, pooled cells*: the test of Meng, Rosenthal and Rubin (1992) for two correlations sharing one variable, on all shared genotype–environment cells after centring observations and both predictions within each environment, so that cells are treated as independent. *Meng within environments, combined*: the same test in each environment, combined by Stouffer's method. *t test over environments*: the per-environment differences in Pearson $r$, with environments as the units. The last rule is the threshold of Section 3.3 (common bean 0.066, spring wheat 0.213, soybean 0.116); because the threshold is fitted to these pairs, its row is in-sample. Table S31 sets the smallest gap a pooled Meng test declares against $3/\sqrt{N}$ in the Gaussian model. Script: analysis/supplementary/code/meng_vs_threshold.py.

***Table S30. Pairs declared different, and whether the more accurate method selects better.***

| Dataset | Rule | Pairs | Declared different | Share declared | More accurate selects better, if declared | Declared pairs at or above the threshold | More accurate selects better, if not declared |
|---|---|---|---|---|---|---|---|
| common bean | Meng, pooled cells | 133 | 74 | 56 % | 80 % | 5 % | 63 % |
| common bean | Meng within | 133 | 52 | 39 % | 83 % | 8 % | 65 % |

| | | | | | | | |
|---|---|---|---|---|---|---|---|
| | environments, combined | | | | | | |
| common bean | t test over environments | 133 | 55 | 41 % | 85 % | 7 % | 63 % |
| common bean | gap at or above the threshold | 133 | 4 | 3 % | 100 % | 100 % | 71 % |
| spring wheat | Meng, pooled cells | 132 | 107 | 81 % | 83 % | 30 % | 68 % |
| spring wheat | Meng within environments, combined | 132 | 109 | 83 % | 83 % | 29 % | 65 % |
| spring wheat | t test over environments | 132 | 109 | 83 % | 84 % | 29 % | 61 % |
| spring wheat | gap at or above the threshold | 132 | 32 | 24 % | 100 % | 100 % | 74 % |
| soybean | Meng, pooled cells | 324 | 299 | 92 % | 80 % | 24 % | 76 % |
| soybean | Meng within environments, combined | 324 | 297 | 92 % | 80 % | 24 % | 70 % |
| soybean | t test over environments | 324 | 244 | 75 % | 86 % | 29 % | 59 % |
| soybean | gap at or above the threshold | 324 | 71 | 22 % | 96 % | 100 % | 75 % |

***Table S31. Smallest accuracy gap a pooled Meng test declares, against 3/√N.***

Gaussian model, mean accuracy 0.3, two-sided 0.05; $r_{12}$ is the correlation between the two methods' predictions. 3/√*N* is the gap at which the more accurate method selects better with probability 0.95 under the design grid of Table S11.

| **Cells *N*** | **Meng, $r_{12}$ = 0.5** | **Meng, $r_{12}$ = 0.8** | **Meng, $r_{12}$ = 0.95** | **3/√*N*** |
|---|---|---|---|---|
| 500 | 0.083 | 0.053 | 0.026 | 0.134 |
| 1,000 | 0.058 | 0.037 | 0.019 | 0.095 |
| 2,000 | 0.041 | 0.026 | 0.013 | 0.067 |
| 4,000 | 0.029 | 0.019 | 0.009 | 0.047 |
| 8,000 | 0.021 | 0.013 | 0.007 | 0.034 |
| 16,000 | 0.015 | 0.009 | 0.005 | 0.024 |

### Section S23. Population structure of the constructed panels

For the genotypes each panel analyses, the markers the panel reads were mean-imputed, monomorphic markers dropped, standardised and decomposed by singular value decomposition, as in the panel scripts (Table S32); the realised relationship matrix is VanRaden's first method. The soybean count is that of the panel file (the analyses use 1,161 of these genotypes in the 169 environments with at least 25 genotypes). The G2F competition analysis uses no markers, and the Chinese maize population is described in its source publications (Section S11). Script: analysis/supplementary/code/pop_structure.py.

***Table S32. Marker structure of the three panels.***

| **Dataset** | **Genotypes** | **Markers** | **PC1 (%)** | **PC2 (%)** | **PCs for 50 %** | **PCs for 80 %** | **Off-diagonal VanRaden relationship, median** | **95th percentile** |
|---|---|---|---|---|---|---|---|---|
| common bean | 467 | 5,820 | 9.2 | 5.3 | 19 | 73 | −0.034 | 0.487 |
| spring wheat | 384 | 2,302 | 6.4 | 4.0 | 42 | 137 | 0.067 | 0.511 |
| soybean | 1,162 | 5,158 | 4.2 | 2.8 | 64 | 254 | −0.024 | 0.271 |

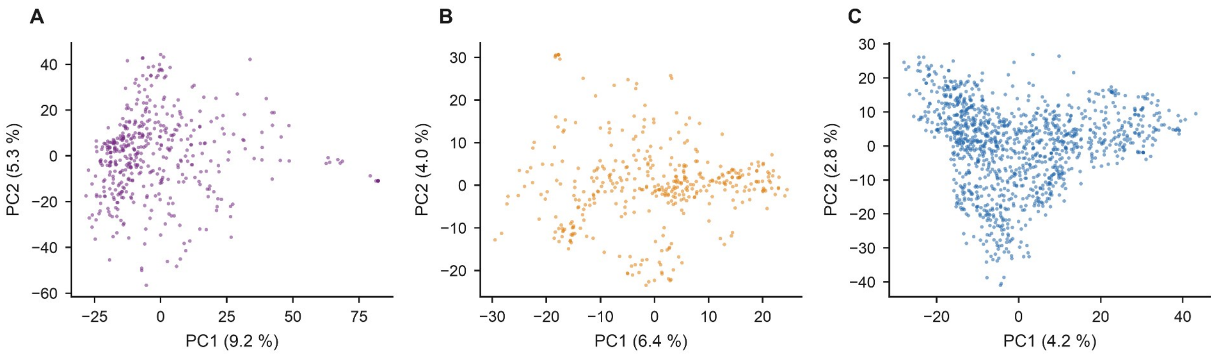

**Fig. S4.** Population structure of the three constructed panels: the first two principal components of the standardised markers, with the share of marker variance each carries. (A) Common bean (VEF Andean elite panel; 467 genotypes, 5,820 markers). (B) Spring wheat (URSN; 384 genotypes, 2,302 markers). (C) Soybean (NUST; 1,162 genotypes, 5,158 markers).

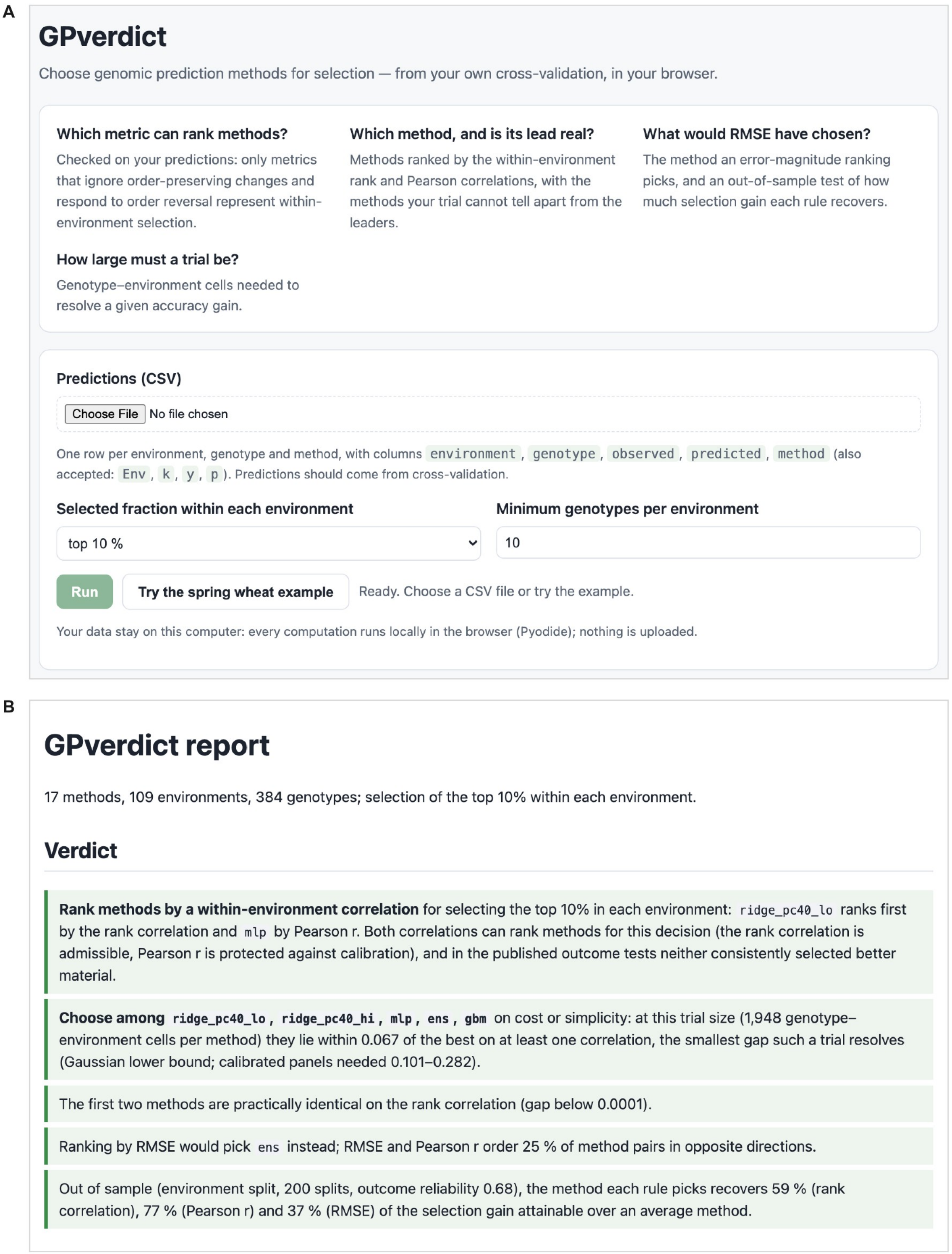

**Fig. S5.** The GPverdict web page (version 1.1.1) running its bundled spring wheat example. (A) The input form; (B) the report it produced, cropped to its verdict section. Unaltered screenshots taken in headless Chrome; the analysis runs locally in the browser (Pyodide) and no data are uploaded.
